# Examining molecular determinants underlying heterogeneity of synaptic release probability using optical quantal imaging

**DOI:** 10.1101/240549

**Authors:** Yulia Akbergenova, Yao V. Zhang, Shirley Weiss-Sharabi, Karen L. Cunningham, J. Troy Littleton

## Abstract

Neurons communicate through neurotransmitter release at specialized synaptic regions known as active zones (AZs). Using transgenic biosensors to image postsynaptic glutamate receptor activation following single vesicle fusion events at *Drosophila* neuromuscular junctions, we analyzed release probability (*P_r_*) maps for a defined connection with ~300 AZs between synaptic partners. Although *P_r_* was very heterogeneous, it represented a stable and unique feature of each AZ. *P_r_* heterogeneity was not abolished in mutants lacking Synaptotagmin 1, suggesting the AZ itself is likely to harbor a key determinant(s). Indeed, AZ *Pr* was strongly correlated with presynaptic Ca^2+^ channel density and Ca^2+^ influx at single release sites. In addition, *P_r_* variability was reflected in the postsynaptic compartment, as high *P_r_* AZs displayed a distinct pattern of glutamate receptor clustering. Developmental analysis suggests that high *P_r_* sites emerge from earlier formed AZs, with a temporal maturation in transmission strength occurring over several days.

## Introduction

Synaptic vesicle fusion occurs at specialized regions of the presynaptic membrane known as active zones (AZs). Several evolutionarily conserved structural proteins are enriched in this subdomain of the presynaptic terminal, including RIM, RIM binding protein, Syd-1, Liprin-α, ELKS/CAST/Bruchpilot, Munc13, and Bassoon/Piccolo/Fife (Schoch and Gundelfinger, 2006; Südhof, 2012; Van Vactor and Sigrist, 2017; Zhai and Bellen, 2004). These large macromolecular complexes facilitate clustering of synaptic vesicles and voltage-gated Ca^2+^ channels (VGCCs). The clustering of VGCCs at AZs allow action potential-triggered Ca^2+^ influx to act locally on synaptic vesicles that are docked and primed for release (Acuna et al., 2016; Bucurenciu et al., 2008; Eggermann et al., 2011; Fouquet et al., 2009; Kawasaki et al., 2004). Synaptic vesicle fusion is tightly regulated and occurs through a highly probabilistic process, often with only a small percent of action potentials triggering release from individual AZs (Körber and Kuner, 2016). Although AZs are thought to share the same overall complement of proteins, release probability (*P_r_*) for synaptic vesicle fusion is highly variable across different neurons, and even across AZs formed by the same neuron (Atwood and Karunanithi, 2002; Branco and Staras, 2009; Melom et al., 2013; Peled and Isacoff, 2011). Indeed, some AZ-specific proteins are non-uniformly distributed, and the molecular composition of AZs can undergo rapid changes (Glebov et al., 2017; Graf et al., 2009; Liu et al., 2016; Reddy-Alla et al., 2017; Sugie et al., 2015; Tang et al., 2016; Weyhersmüller et al., 2011; Wojtowicz et al., 1994).

The *Drosophila* neuromuscular junction (NMJ) has emerged as a useful system to study release heterogeneity. At this connection, motor neurons form glutamatergic synapses onto bodywall muscles in a stereotypical fashion, with the axon expanding to form ~10–60 synaptic boutons that each contain many individual AZs (Harris and Littleton, 2015). *Drosophila* AZs are termed T-bars due to their mushroom-like appearance by EM, and contain a similar assortment of proteins to those identified at mammalian AZs (Böhme et al., 2016; Bruckner et al., 2017, 2012; Ehmann et al., 2014; Feeney et al., 1998; Fouquet et al., 2009; Graf et al., 2012; Jan and Jan, 1976; Kaufmann et al., 2002; Kittel et al., 2006; Liu et al., 2011; Owald et al., 2010; Wagh et al., 2006). Each AZ is specifically associated with a postsynaptic glutamate receptor field, usually separated from other PSDs by membrane infoldings that form the subsynaptic reticulum (SSR) (Johansen et al., 1989). Glutamate receptors at the *Drosophila* NMJ are excitatory inotropic non-NMDA receptors that exist as tetramers, with three obligatory subunits encoded by GluRIII, GluRIID and GluRIIE, and a variable 4^th^ subunit encoded by either GluRIIA (A-type) or GluRIIB (B-type) (Featherstone et al., 2005; Marrus et al., 2004; Petersen et al., 1997; Qin et al., 2005; Schuster et al., 1991). GluRIIA containing receptors generate a larger quantal size and display slower receptor desensitization that their GluRIIB counterparts (DiAntonio et al., 1999). The A- and B-subtypes compete for incorporation into the tetramer at individual postsynaptic densities (PSDs) in a developmental and activity-regulated fashion (Chen and Featherstone, 2005; DiAntonio et al., 1999; Marrus and DiAntonio, 2004; Rasse et al., 2005; Schmid et al., 2008).

The stereotypical alignment of individual AZs to distinct postsynaptic glutamate receptor fields in *Drosophila* allowed the generation of genetic tools to optically follow quantal fusion events at single release sites by visualizing glutamate receptor activation (Melom et al., 2013; Peled and Isacoff, 2011). Classically, studies of synaptic transmission have used electrophysiology to measure the postsynaptic effect of neurotransmitter release over a population of release sites (Katz and Miledi, 1969, 1967), precluding an analysis of how individual AZs respond. By transgenically expressing GCaMP Ca^2+^ sensors that target to the postsynaptic membrane through PDZ binding or myristoylation domains, single vesicle fusion events at each individual AZ can be imaged by following the spatially localized Ca^2+^ influx induced upon glutamate receptor opening. This allows for the generation of *P_r_* maps for both evoked and spontaneous fusion for all AZs formed onto the muscle by the innervating motor neuron (Cho et al., 2015; Melom et al., 2013; Muhammad et al., 2015; Newman et al., 2017; Peled et al., 2014; Peled and Isacoff, 2011; Reddy-Alla et al., 2017). One surprising finding using this quantal imaging approach is that the hundred of AZs formed by a single motor neuron have a heterogeneous distribution of *P_r_*, ranging from 0.01 to ~ 0.5, with neighboring AZs often showing greater than 40-fold differences in *P_r_* (Melom et al., 2013; Peled et al., 2014).

Key questions raised by these observations include how *P_r_* is uniquely set for individual AZs, and whether rapid changes in *P_r_* mediate distinct forms of synaptic plasticity. One potential mechanism to explain *P_r_* variability is that presynaptic AZs show distinct Ca^2+^ channel density and subsequent Ca^2+^ influx at single release sites. An alternative model is that Ca^2+^ entry is similar over all AZs, and that evoked *P_r_* is correlated with local cytosolic or synaptic vesicle proteins and their number and/or state (i.e. phosphorylation status). Likewise, another AZ determinant beyond Ca^2+^ channels could be differentially distributed that controls synaptic vesicle docking or priming state. Each of these possibilities could contribute to AZ *P_r_* collectively, or one mechanism might dominate. To gain insight into the molecular mechanisms that control the heterogeneous distribution of *P_r_* at AZs, we employed optical quantal imaging at *Drosophila* NMJs to identify high *P_r_* sites and examine their properties. Our findings indicate that AZs show differential accumulation of Ca^2+^ channels that generate distinct levels of presynaptic Ca^2+^ influx and variable *P_r_*, with AZ maturation playing a key role in setting *Pr*.

## Results

### *Drosophila* NMJ synapses display heterogeneity in *P_r_*, ranging from functionally silent sites to high *P_r_* AZs

Recent studies have demonstrated that release sites possess unique structural and functional heterogeneity (Éltes et al., 2017; Holderith et al., 2012; Maschi and Klyachko, 2017; Melom et al., 2013; Peled et al., 2014; Reddy-Alla et al., 2017; Sugie et al., 2015). Using the *Drosophila* NMJ, we explored the source of variation in *P_r_* at this synaptic connection. We previously observed that evoked *P_r_* is non-uniform across a population of ~300 AZs formed by motor neuron MN4-Ib onto muscle 4, ranging from 0.01 to ~0.5 in HL3 saline containing 1.3 mM extracellular Ca^2+^ and 20 mM Mg^2+^ (Melom et al., 2013). In our original study, each AZ was defined by ROIs where postsynaptic Ca^2+^ flashes were observed during stimulation, but AZs were not co-labeled in the live preparation. To more closely examine AZ *P_r_* heterogeneity, we identified the position of each corresponding PSD by co-expressing the RFP-tagged glutamate receptor subunit GluRIIA (Rasse et al., 2005). We also created a newer edition of our previous biosensor by generating UAS transgenic animals expressing the Ca^2+^ sensitive GCaMP6s fused with an N-terminal myristoylation (myr) domain. We targeted UAS-myrGCaMP6s to the postsynaptic muscle membrane using Mef2-GAL4 and monitored Ca^2+^ influx after activation of postsynaptic glutamate receptors from spontaneous neurotransmitter release or nerve stimulation (0.3 Hz for 5 minutes) in muscle 4 of early stage 3^rd^ instar larvae (Movie 1). In response to stimulation, synchronous vesicle fusion could be identified across distinct populations of release sites (Figure 1A). During multiple rounds of stimulation, fusion events corresponding to release from different subsets of AZs were observed. However, a small subset of AZs (~10%) experienced far more frequent fusion events than average, with GCaMP6s activation at the same PSD repeatedly observed during low frequency stimulation (Figure 1A, arrow).

**Figure 1.**
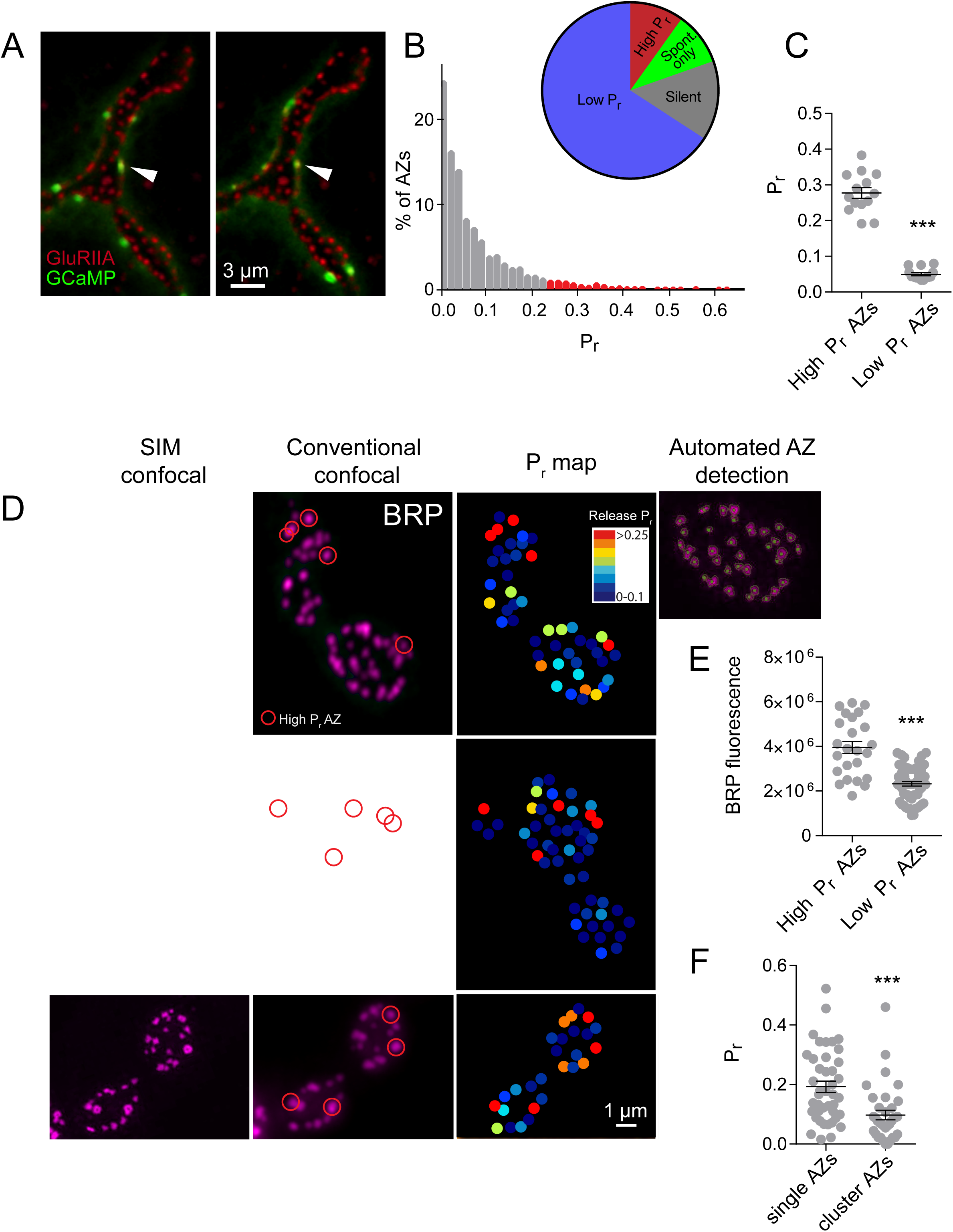
High *P_r_* sites correspond to single AZs with elevated levels of BRP. (**A**) Representative images of consecutive evoked release events (green flashes) visualized by expressing myrGCaMP6s in muscle 4. The position of each AZ was determined by expressing GluRIIA-RFP to label the corresponding PSD. Evoked release triggers fusion across different sets of AZs during each stimuli, but a subpopulation of AZs respond more frequently (arrow). (**B**) Histogram of the distribution of AZ *P_r_* for a 0.3 Hz 5-minute stimulation paradigm. AZs classified as high *P_r_* (>2 standard deviations above the mean) are shown in red. The percentage of AZs that were low *P_r_* (65.8%), high *P_r_* (9.9%), spontaneous-only (9.7%) and silent (14.6%) is displayed in the inset. (**C**) Average *P_r_* determined for each individual experiment for the AZ population categorized based on low and high activity sites (>2 standard deviations above the mean). Each point represents the average for all AZs (classified as either high or low *Pr*) from a single animal. (**D**) Individual BRP puncta for three NMJs from three different animals imaged with high resolution structured illumination microscopy (SIM, left panel) or confocal microscopy (middle panel). The right panel displays the heat map for evoked *P_r_* from the same NMJs determined by GCaMP6s imaging prior to fixation. Representative high *P_r_* sites are circled with red in the middle panels. The far right top panel displays the results from the automated detection algorithm that outlines individual AZs. (**E**) AZs were separated into high and low *P_r_* based on their activity and the fluorescence intensity of the corresponding BRP puncta is shown (from conventional confocal images). (**F**) AZs with high BRP intensity (two standard deviations above average) were preselected from conventional confocal images and identified on corresponding SIM images. In cases where the BRP signal was resolvable into more than one AZ by SIM microscopy, it was assigned to the AZ cluster group. In cases where the BRP signal mapped to a single BRP puncta by SIM imaging, it was assigned to the single AZ group. *P_r_* is plotted for each group. Student’s t-test was used for statistical analysis (*** = p≤0.001). Error bars represent SEM.

Using this approach, we mapped all myrGCaMP6s visualized release events to the actual position of *in vivo* labeled glutamate receptor PSDs marked by GluRIIA-RFP. Consistent with our previous data, we observed a heterogeneous distribution of AZ *P_r_*, with an average *P_r_* of 0.073 ± 0.004 (n = 1933 AZs from 16 NMJs from 16 animals). However, there was a >50-fold difference in *P_r_* between the highest and lowest releasing sites. We plotted the distribution of AZ *P_r_* and observed a skew in the data, with a small number of AZs consistently showing high release rates (75% percentile of *P_r_* was 0.1, with a maximum *P_r_* of 0.73, Figure 1B). Indeed, the release probability data did not fit a normal distribution (D'Agostino K^2^ test (p<0.0001), Shapiro-Wilk test (p<0.0001), Kolmogorov-Smirnov test (p<0.0001)). Beyond the heterogeneous *P_r_* distribution, 9.7% of all release sites with apposed GluRIIA receptors displayed only spontaneous fusion events, and another 14.6% of the AZ population was silent for both spontaneous and evoked release during the recording period (Figure 1B). In addition, the majority of AZs rarely released a synaptic vesicle following an action potential, with a *P_r_* in the range of 0.01 to 0.2. To functionally examine differences between high and low releasing sites, we categorized all AZs with a release rate greater than 2 standard deviations above average as “high *P_r_*”, and the remaining AZs that showed evoked release as “low *P_r_*”. Using these criteria, 65.8% of all AZs fell in the low *P_r_* category with an average release probability of 0.049 ± 0.004. 9.9% of AZs were classified as high *P_r_* sites, with an average release probability of 0.277 ± 0.015 (Figure 1C), indicating high *P_r_* AZs displayed on average a 5.7-fold increased chance of vesicle fusion following an action potential compared to low *P_r_* AZs.

### High *P_r_* AZs correspond to single release sites with enhanced levels of the AZ protein Bruchpilot

One potential caveat to the interpretation of heterogeneous distributions in *P_r_* is the possibility that multiple release sites positioned in proximity to each other contribute to a false identification of high *P_r_* sites. To determine if high *P_r_* sites were due to release from multiple neighboring AZs, we employed high-resolution structured illumination microscopy (SIM) (Figure 1D) and combined it with the quantal imaging method. Presynaptic AZ position can be precisely identified at the NMJ by labeling the core T-bar component Bruchpilot (BRP), the homolog of mammalian ELKS/CAST (Fouquet et al., 2009; Wagh et al., 2006). Using dual color imaging (myrGCaMP6s and GluRIIA-RFP), we first mapped *P_r_* across all AZs, and then fixed the tissue and labeled with anti-BRP antisera. SIM microscopy provides a lateral resolution of labeled biological structures with a limit of ~110 nm (Wegel et al., 2016), allowing separation of individual AZs where the average BRP ring diameter is ~200 nm (Owald et al., 2012). The presence of GluRIIA-RFP allowed precise mapping of release sites between live and fixed tissue, as well as correlation of high *P_r_* sites with SIM labeled BRP-positive AZs (Figure 1D). Using an automated detection algorithm in the Volocity 3D image analysis software, we were able to identify all AZs labeled with BRP (Figure 1D, right panel), and to resolve individual AZ clusters that were not separated using conventional spinning disk microscopy where *P_r_* was determined. The theoretical lateral resolution of the spinning disk confocal microscope for RFP labeled structures is ~280 nm. Analysis of distances between different AZs by SIM indicated that 2.45 ± 0.4% (n = 9 NMJs from 9 animals) of all AZs were located close enough to each other (within 280 nm) such that they would not be resolvable in our live imaging. In contrast, 9.9% (n = 16 NMJs from 16 animals) of AZs were functionally classified as high *P_r_* sites. Therefore, the majority of high releasing sites are not likely to be explained by release events occurring from closely linked AZs.

To further analyze single versus closely spaced AZs, release maps were generated where *P_r_* was color-coded to visualize the range of probabilities for all release sites. *P_r_* maps were then compared to BRP-positive AZs identified by SIM. As shown in Figure 1D, the vast majority of high *P_r_* sites were represented by a single BRP-positive AZ that was not further resolvable after SIM imaging (Figure 1D, red circles). These single BRP clusters at high *P_r_* sites were larger and brighter than most other BRP positive puncta (Figure 1E). The average total fluorescence of single BRP puncta from high *P_r_* AZs (3.95 × 10^6^ ± 2.67 × 10^5^, n = 24 AZs from 9 NMJs from 9 animals) was 1.7-fold greater than the fluorescence of randomly selected low *P_r_* BRP clusters (2.33 × 10^6^ ± 0.98 × 10^5^, n = 60 AZs from 9 NMJs from 9 animals, p<0.0001). To further examine these large single BRP clusters and their release properties, larger clusters that could not be resolved using conventional spinning disk microscopy were separately analyzed. All BRP clusters larger than 280 nm were automatically detected and assigned their release probability parameters measured during live imaging. We then determined whether these sites were represented by single or multiple AZs using SIM microscopy. Clusters > 280 nm in diameter that could be resolved to multiple BRP positive AZs after SIM imaging had a lower *P_r_* (0.10 ± 0.02, n = 35 AZs from 5 NMJs from 5 animals) than those comprised of a single large BRP positive AZ (0.19 ± 0.02, n = 42 AZs from 5 NMJs from 5 animals, Figure 1F). As such, high resolution SIM microscopy confirms that most high *P_r_* sites correspond to single AZs with more intense BRP labeling, consistent with previous data regarding the positive role of BRP in regulating *P_r_* (Peled et al., 2014; Reddy-Alla et al., 2017).

### Individual AZ *P_r_* is stable across imaging sessions

We next examined if the non-uniform distribution of *P_r_* across AZs was stable when no plasticity changes were induced. If *P_r_* was highly dynamic at individual AZs over time, unique local synaptic vesicle pools might be an important contributor to the distribution of variable release properties. However, a more stable *P_r_* would argue for a specific factor(s) resident at individual AZs. We were limited in our ability to examine *P_r_* continuously over time intervals greater than 10–15 minutes due to bleaching of GCaMP6s from the high frequency capture rate. Within this constraint, we conducted a 3-minute imaging session using 0.3 Hz stimulation to generate an initial *P_r_* map, and then allowed the preparation to rest for 5 minutes without stimulation or imaging. *P_r_* was then re-mapped in a final 3-minute imaging session using 0.3 Hz stimulation. The activity level of individual AZs was very stable between the two sessions (Figure 2A). This was especially evident for high *P_r_* sites, which sustained high levels of activity during both imaging sessions. Plotting the release rate for all AZs revealed a strong correlation for *P_r_* across the two imaging sessions (Pearson r = 0.77, R^2^=0.59, p<0.0001, n = 988 AZs from 8 NMJs from 7 animals, Figure 2B). These data suggest that release rate is a unique property of each AZ and is stable over this time interval.

**Figure 2.**
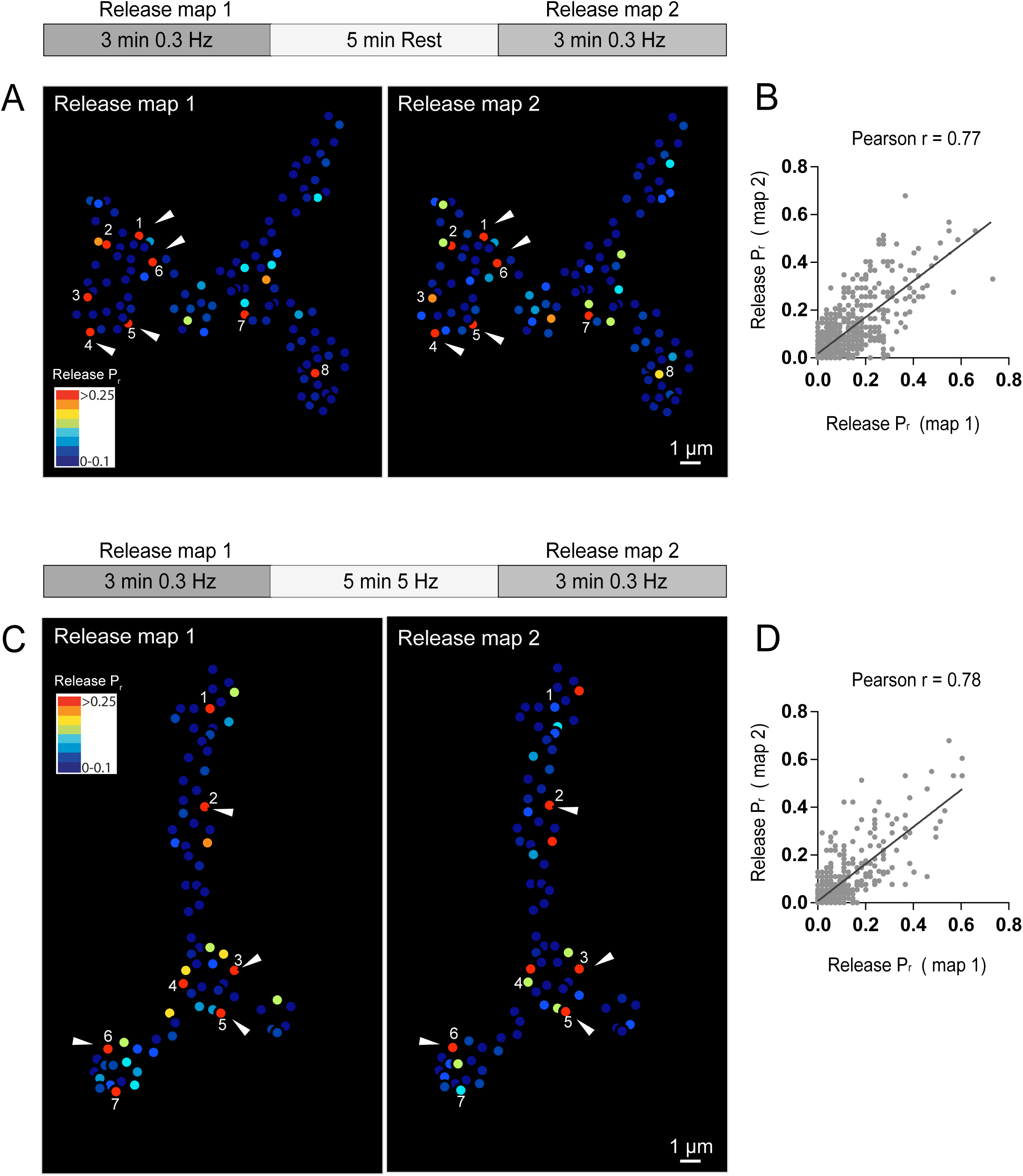
Stability of release maps at the NMJ. (**A**) *P_r_* heatmaps for the same muscle 4 NMJ were generated for two individual imaging sessions, separated by a 5-minute resting period. High *P_r_* AZs were numbered and re-identified in each heatmap. Representative high *P_r_* AZs that sustain release rates during the second imaging session are noted with arrows. (**B**) Correlation of AZ *P_r_* between two imaging sessions separated by a 5-minute resting period. (**C**) *P_r_* heatmaps for the same NMJ separated by a 5-minute 5 Hz stimulation. Representative high *P_r_* AZs that did not change activity levels are noted with arrows. (**D**) Correlation of AZ *P_r_* between two imaging sessions separated by a 5-minute 5 Hz stimulation period.

Heterogeneous release rates between AZs might be sensitive to the accumulation of different vesicle populations with variable levels of fusogenicity. If so, a stronger stimulation paradigm that is sufficient to drive vesicle cycling and intermixing would be expected to alter the *P_r_* map. To test this, NMJ preparations were imaged during two low frequency 0.3 Hz stimulation periods separated by a 5-minute 5 Hz stimulation session to induce robust synaptic vesicle turnover and recycling (Figure 2C). Release maps were not dramatically altered by 5 Hz stimulation, with the overall correlation of *P_r_* similar to maps generated without stimulation (Pearson r = 0.78, R^2^ = 0.61, p<0.0001, n = 613 AZs from 6 NMJs from 6 animals, Figure 2D). Thus, inducing vesicle recycling with 5 Hz stimulation does not dramatically change *P_r_* across the AZ population, arguing that structural AZ components, versus specific vesicle populations surrounding AZs, are likely to represent the major driver of *P_r_* heterogeneity at this synapse.

### *Synaptotagmin* null mutants display release heterogeneity across AZs

The synchronous Ca^2+^ sensor Synaptotagmin 1 (Syt1) resides on synaptic vesicles and plays a major role in *P_r_* determination at *Drosophila* NMJs (DiAntonio and Schwarz, 1994; Guan et al., 2017; Lee et al., 2013; Littleton et al., 1994, 1993; Yoshihara et al., 2003; Yoshihara and Littleton, 2002). We hypothesized that if synaptic vesicle proteins play a key role in *P_r_* heterogeneity, in addition to their established role in determining overall *P_r_*, then elimination of Syt1 would likely disrupt this heterogeneity. We therefore expressed myrGCaMP6s with Mef2-GAL4 in *syt1* null mutants. As observed electrophysiologically, quantal imaging in *syt1* null mutants revealed a dramatic reduction in evoked release, a shift from synchronous to highly asynchronous fusion, and an increase in spontaneous release rates (Movie 2). To estimate AZ heterogeneity in *syt1* nulls, preparations were stimulated at 5 Hz and release events were normalized to the number of stimuli (Figure 3A). The average release rate per AZ per second in *syt1* nulls during 5 Hz stimulation was 0.03 ± 0.001 (n = 719 AZs from 7 NMJs from 6 animals, Figure 3B). In contrast, spontaneous release rate per AZ in the absence of stimulation was 0.018 ± 0.001 per second in *syt1* nulls (n = 719 AZs from 7 NMJs from 6 animals) compared to 0.011 ± 0.001 in controls (n = 559 AZs from 6 NMJs from 4 animals, p<0.0001, Figure 3B). All visualized release events were mapped to specific AZs and representative *P_r_* heatmaps were generated (Figure 3A). Although release rate is dramatically reduced in *syt1* nulls, AZs still maintain the overall heterogeneity in *P_r_* distribution across AZs (Figure 3C-E). Comparing the distribution of AZ release rates for *syt1* nulls and controls, release was proportionally decreased across all AZs in *syt1* (Figure 3C). Frequency distribution analysis of AZs with normalized release rates (from 0 to maximum release) confirmed that there was no significant change in the heterogeneity of release between *syt1* mutants and controls (Figure 3D). Likewise, the cumulative frequency distribution of normalized AZ *P_r_* was similar between *syt1* mutants and controls (Figure 3E). Given that AZ release remains highly heterogeneous in the absence of Syt1, these data suggest that variable distribution of key AZ components, rather than heterogeneity of local synaptic vesicle proteins, is likely to control *P_r_* distribution across *Drosophila* NMJ AZs.

**Figure 3.**
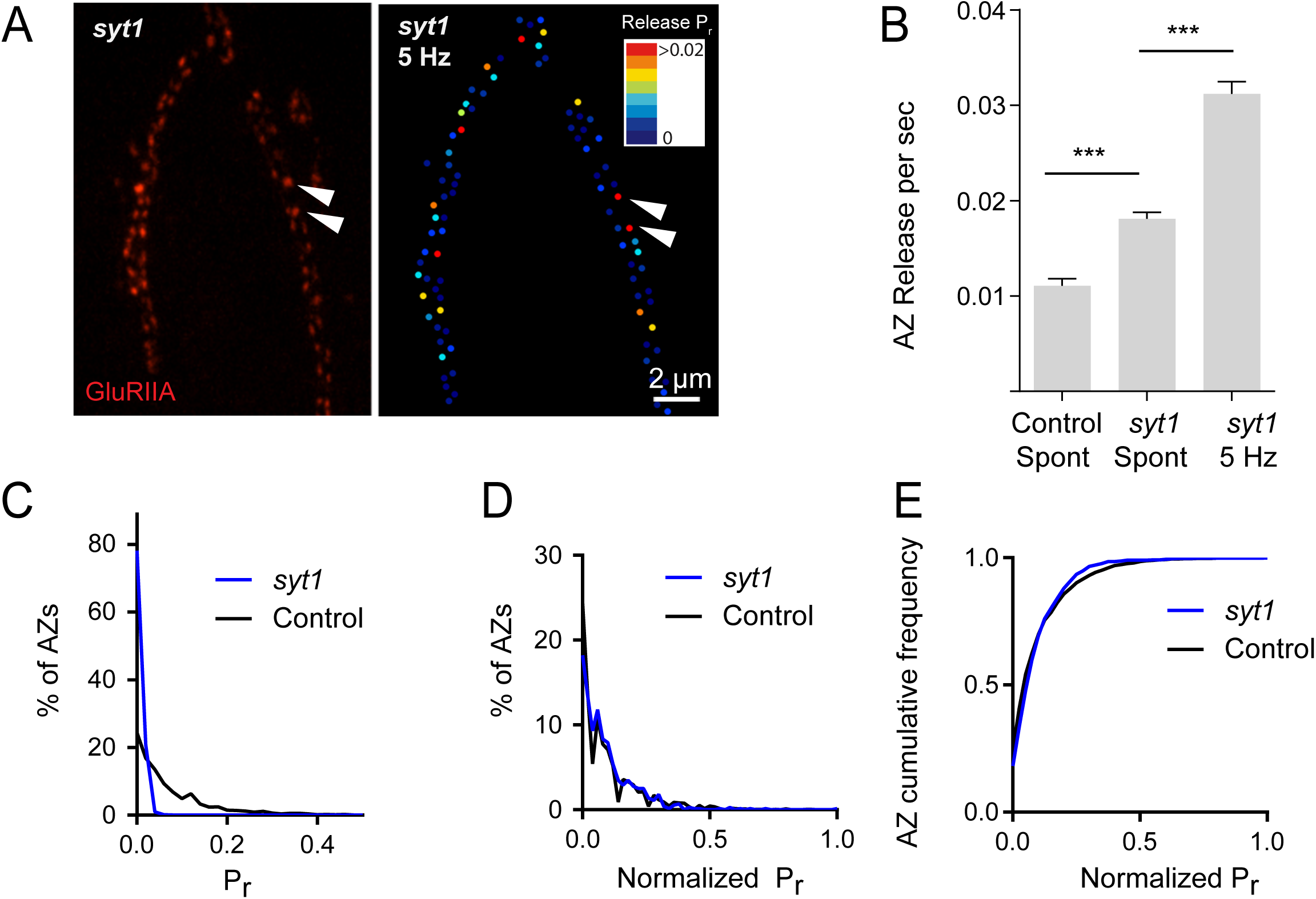
*P_r_* variability remains in *syt1* null mutants. (**A**) The left panel displays the distribution of GluRIIA staining in *syt1* nulls (left panel) at the muscle 4 NMJ. The corresponding *P_r_* heatmap is shown on the right. The arrows denote several high *P_r_* sites opposed by bright GluRIIA positive PSDs. (**B**) AZ release events per second for spontaneous release and evoked by 5 Hz stimulation are shown for *syt1* nulls mutants, and for spontaneous release in controls. (**C**) Frequency distribution of *P_r_* is shown for *syt1* nulls and controls. (**D**) Plot of normalized *P_r_* frequency distribution (from 0 to 1 (max)) for *syt1* nulls and controls. (**E**) Cumulative frequency distribution for normalized release rates for *syt1* nulls and controls is shown. Student’s t-test was used for statistical analysis (*** = p≤0.001). Error bars represent SEM.

### *P_r_* is highly correlated with Ca^2+^ channel abundance at AZs

We next investigated what key AZ protein(s) might control *P_r_*. Many structural components of AZs cooperate to regulate positioning of synaptic vesicles in the vicinity of Ca^2+^ channels. Indeed, synaptic vesicle fusion is highly sensitive to Ca^2+^ and most effective in close proximity to Ca^2+^ channels (Augustine et al., 1985; Böhme et al., 2016; Chen et al., 2015; Heidelberger et al., 1994; Katz and Miledi, 1967; Katz, 1969; Keller et al., 2015; Meinrenken et al., 2003, 2002; Stanley, 2016; Wang et al., 2008). As such, Ca^2+^ channel abundance and the subsequent level of Ca^2+^ influx at individual AZs is a compelling variable for *P_r_* heterogeneity. In *Drosophila*, Cacophony (*cac*) encodes the voltage-activated Ca^2+^ channel a1 subunit required for neurotransmitter release (Fouquet et al., 2009; Kawasaki et al., 2004, 2000; Littleton and Ganetzky, 2000; Liu et al., 2011; Rieckhof et al., 2003; Smith et al., 1996). Transgenic animals expressing fluorescently tagged Cac channels have been previously generated, demonstrating that Cac localizes specifically to AZs at the NMJ (Kawasaki et al., 2004; Matkovic et al., 2013; Yu et al., 2011). To examine the effect of differential Cac distribution on AZ release, dual color imaging experiments were performed where vesicle fusion events were detected by myrGCaMP6s and Ca^2+^ channel distribution was visualized by expression of red-labeled Cac-TdTomato. Preparations were stimulated at 0.3 Hz for 5 minutes and AZ release rate was compared with Cac-TdTomato fluorescence distribution (Figure 4A, B). A strong positive correlation (Pearson r = 0.62, R^2^ = 0.38, p<0.0001, n = 483 AZs from 7 NMJs from 7 animals) was observed between Cac fluorescence intensity and evoked AZ *P_r_* (representative experiment shown in Figure 4B). We next examined if there was a similar correlation between Cac levels and the frequency of spontaneous vesicle release (minis) at individual AZs. AZ release rates for spontaneous events showed only a mild correlation (Pearson r = 0.19, R^2^ = 0.04, p<0.0001, n = 483 AZs from 7 NMJs from 7 animals) between mini frequency and Cac density (representative experiment shown in Figure 4C). These results match well with previous observations that release rates for evoked and spontaneous fusion are poorly correlated at *Drosophila* AZs (Melom et al., 2013; Peled et al., 2014), and that spontaneous fusion is largely independent of extracellular Ca^2+^ at this synapse (Jorquera et al., 2012; Lee et al., 2013).

**Figure 4.**
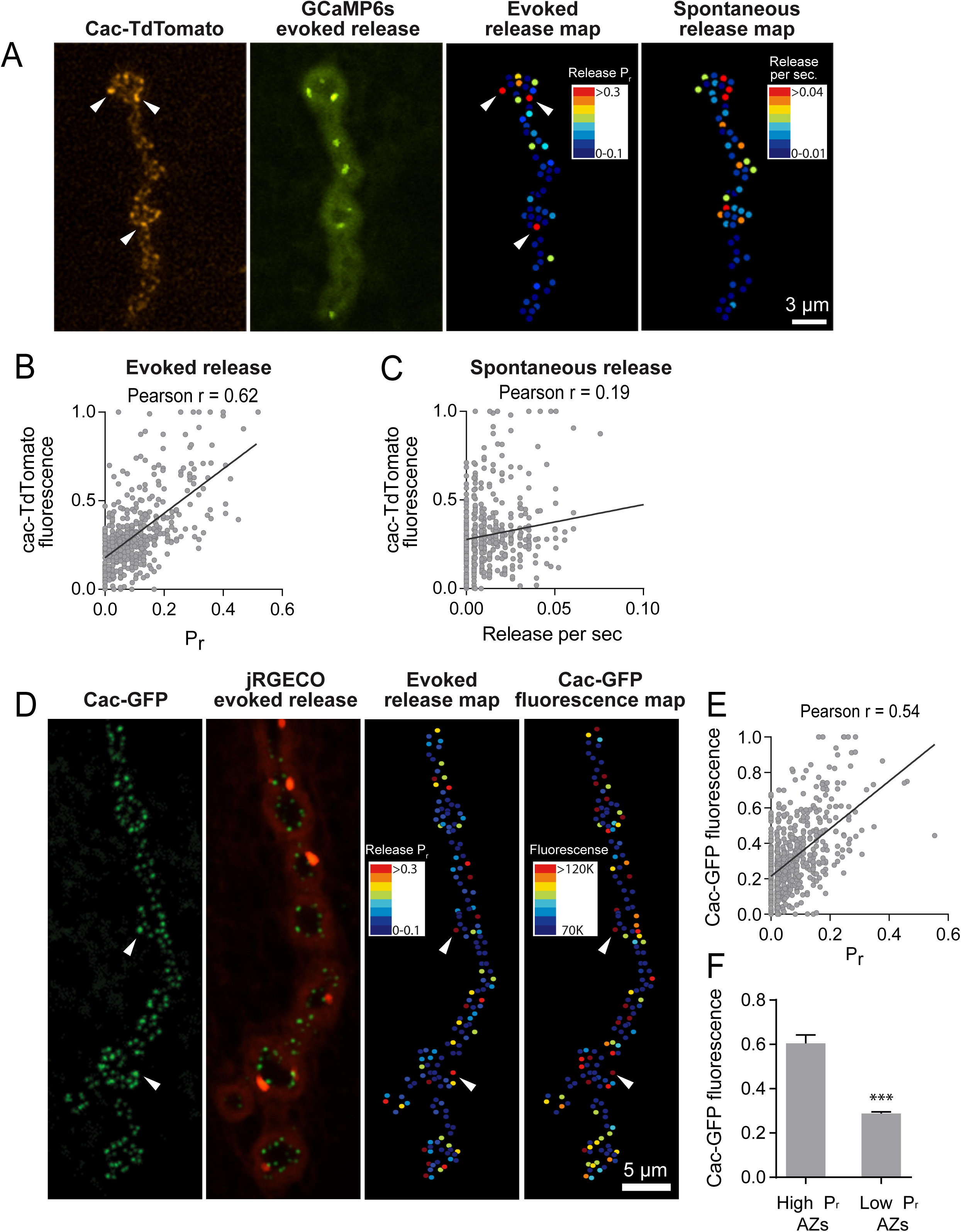
*P_r_* correlates with Cac channel density at AZs. (**A**) Representative images showing heterogeneous distribution of Cac-TdTomato at the NMJ of muscle 4 (left panel). Evoked release was visualized at the same NMJ using myrGCaMP6s (second panel) and AZ release maps were generated for evoked (third panel) and spontaneous fusion (right panel). Several high *P_r_* AZs with bright Cac density are noted (arrows). (**B**) Correlation between AZ *P_r_* and Cac-TdTomato fluorescent intensity for evoked release. (**C**). Correlation between AZ spontaneous release rate per second and Cac-TdTomato fluorescence intensity. (**D**) Representative images showing heterogeneous distribution of Cac-GFP at the NMJ (left panel). Evoked release visualized at the same NMJ by myr-jRGECO is shown in the second panel. The *P_r_* heatmap for evoked release is shown in the third panel. A heatmap distribution of Cac-GFP fluorescence intensities, based on same criteria as color-coding of *P_r_*, is shown in the right panel. The arrows denote several higher *P_r_* sites containing bright Cac-GFP puncta. (**E**) Correlation between AZ *P_r_* and Cac-GFP fluorescence intensity for evoked release. (**F**) Cac-GFP fluorescence obtained for AZs functionally classified as either low or high *P_r_* (>2 standard deviations above mean) by quantal imaging with myr-jRGECO1a. Student’s t-test was used for statistical analysis (*** = p≤0.001). Error bars represent SEM.

To gain confidence that the observed Cac-TdTomato intensity accurately reflects Cac channel distribution, Cac channels transgenically tagged with GFP were also examined. Postsynaptic expression of myrGCaMP6s obscured Cac-GFP fluorescence, preventing quantal imaging with this sensor in the Cac-GFP background. Therefore, transgenic lines expressing myristoylated red Ca^2+^ indicators previously characterized in the field were generated, including RCaMP1h, R-GECO1 and jRGECO1a. Although RCaMP1h and R-GECO1 were too dim to visualize localized Ca^2+^ transients at AZs, transgenic lines expressing the myristoylated Ca^2+^ indicator jRGECO1a (Dana et al., 2016) in muscle 4 allowed detection of Ca^2+^ influx events following vesicle fusion at single AZs (Movie 3). In contrast to the more robust GCaMP6s, jRGECO1a has a shorter fluorescent lifetime and the signal amplitude decays more rapidly over time. Indeed, quantal events imaged in transgenic animals expressing myr-jRGECO1a were dimmer and fully bleached within 7–10 minutes of imaging. Therefore, preparations were stimulated at 1 Hz for shorter two minute imaging sessions to generate *P_r_* maps in myr-jRGECO1a expressing larvae (Figure 4D). Using this approach, a strong correlation (Pearson r = 0.54, R^2^ = 0.29, p<0.0001, n = 651 AZs from 7 NMJs from 7 animals) between AZ *P_r_* detected by myr-jRGECO1a and Cac-GFP density was observed (representative experiment shown in Figure 4E). Again, a weaker correlation was found between rates of spontaneous events and Cac-GFP density (Pearson r = 0.17, R^2^ = 0.03, p<0.0001, n = 651 AZs from 6 NMJs from 6 animals). Hence, *P_r_* for action-potential evoked fusion is strongly correlated with the local density of Cac channels at individual AZs, regardless of which fluorophore is used to visualize Cac.

To determine the relative levels of Cac that defined low and high *P_r_* sites, the distribution of Cac-GFP and BRP across the AZ population was examined using SIM microscopy. Similar to the variable levels of BRP described earlier (Figure 1D), variability in the distribution and mean intensity of Cac-GFP fluorescence across AZs at muscle 4 was observed by SIM (Figure 4 – figure supplement 1A, B). 5.72% of AZs displayed Cac-GFP fluorescence greater than 2 standard deviations above average (n = 2011 AZs from 11 NMJs from 3 animals). The mean Cac-GFP fluorescence for these bright AZs (>2 standard deviations above average) was 2.1-fold greater than that observed for the remaining sites (p<0.0001, Figure 4 – figure supplement 1C). We next compared Cac-GFP fluorescence obtained for AZs that were functionally classified as either low or high *P_r_* sites by quantal imaging using myr-jRGECO1a (Figure 4F). The average fluorescence of single Cac-GFP puncta from high *P_r_* AZs (normalized intensity = 0.6 ± 0.04, n = 38 AZs from 7 NMJs from 7 animals) was 2.09-fold greater than the average fluorescence of low *P_r_* AZs (normalized intensity = 0.29 ± 0.01, n = 638 AZs from 7 NMJs from 7 animals, p<0.0001). The *P_r_* of AZs classified on the basis of the levels of their Cac-GFP fluorescence was also examined. The average *P_r_* for AZs displaying high Cac-GFP fluorescence (>2 standard deviations above average) was 0.2 ± 0.016 (n = 7 NMJs from 7 animals) compared to 0.06 ± 0.003 (n = 7 NMJs from 7 animals, p<0.0001) for the remaining AZs with lower levels of Cac-GFP. Although the absolute number of Cac channels at single AZs is unknown, these data indicate a ~2-fold increase in Ca^2+^ channel number is likely sufficient to change a low *P_r_* site into a high *P_r_* AZ at the *Drosophila* NMJ. Given the steep 3^rd^ to 4^th^ order non-linear dependence of synaptic vesicle fusion with Ca^2+^ (Dodge and Rahamimoff, 1967; Heidelberger et al., 1994; Jan and Jan, 1976), a small change in channel number is likely to have a large effect on *P_r_*.

### *P_r_* correlates with the level of presynaptic Ca^2+^ influx measured at individual AZs

Although our data indicate that Cac channel density correlates with AZ *P_r_*, a more important functional readout of Ca^2+^ channel activity is the local Ca^2+^ influx occurring at each AZ that drives synaptic vesicle fusion. It is unclear if transgenically tagged Ca^2+^ channel fluorescence intensity functionally reflects a heterogeneous level of Ca^2+^ influx at each AZ. Indeed, Ca^2+^ channels are a source of widespread modulation by second messenger pathways that can alter channel conductivity (Catterall and Few, 2008; Dolphin et al., 1991; Evans and Zamponi, 2006; Reid et al., 2003; Tedford and Zamponi, 2006; Zamponi and Snutch, 1998), indicating the abundance of Ca^2+^ channels may not be the best proxy for AZ Ca^2+^ entry. In addition, a direct measure of Ca^2+^ influx would be useful to bypass any potential unknown effects on *P_r_* generated by expressing fluorescently tagged Cac. To generate an estimate of the Ca^2+^ influx that each AZ experiences independent of tagging Cac channels, Ca^2+^ influx was visualized by positioning GCaMP6m near Ca^2+^ channels at the AZ. Cac channels cluster in proximity to the AZ structural protein BRP (Fouquet et al., 2009). The C-terminal fragment of BRP is located further away from the AZ, where it spreads into a filamentous “umbrella” like structure. As such, GCaMP6m was added to the N-terminus of BRP, which localizes directly at the base of the AZ where Ca^2+^ channels cluster. Transgenic animals expressing N-terminal GCaMP6m fusions to a BRP fragment (BRP-short) corresponding to amino acids 473–1226 of the full 1740 amino acid protein (Schmid et al., 2008) were generated. BRP-short targets to AZs and co-localizes with native BRP (Fouquet et al., 2009; Schmid et al., 2008). Expression of GCaMP6m labeled BRP-short did not alter its localization, and GCaMP6m fluorescence could be detected presynaptically at individual AZs at 3^rd^ instar larval NMJs (Figure 5A). At rest, N-terminal ^GCaMP^BRP-short was dim, consistent with the low levels of resting Ca^2+^ inside terminals (Figure 5A). However, stimulation at 10 Hz resulted in a robust increase in discrete punctated fluorescence that remained confined to single AZs during stimulation (Figure 5A).

**Figure 5.**
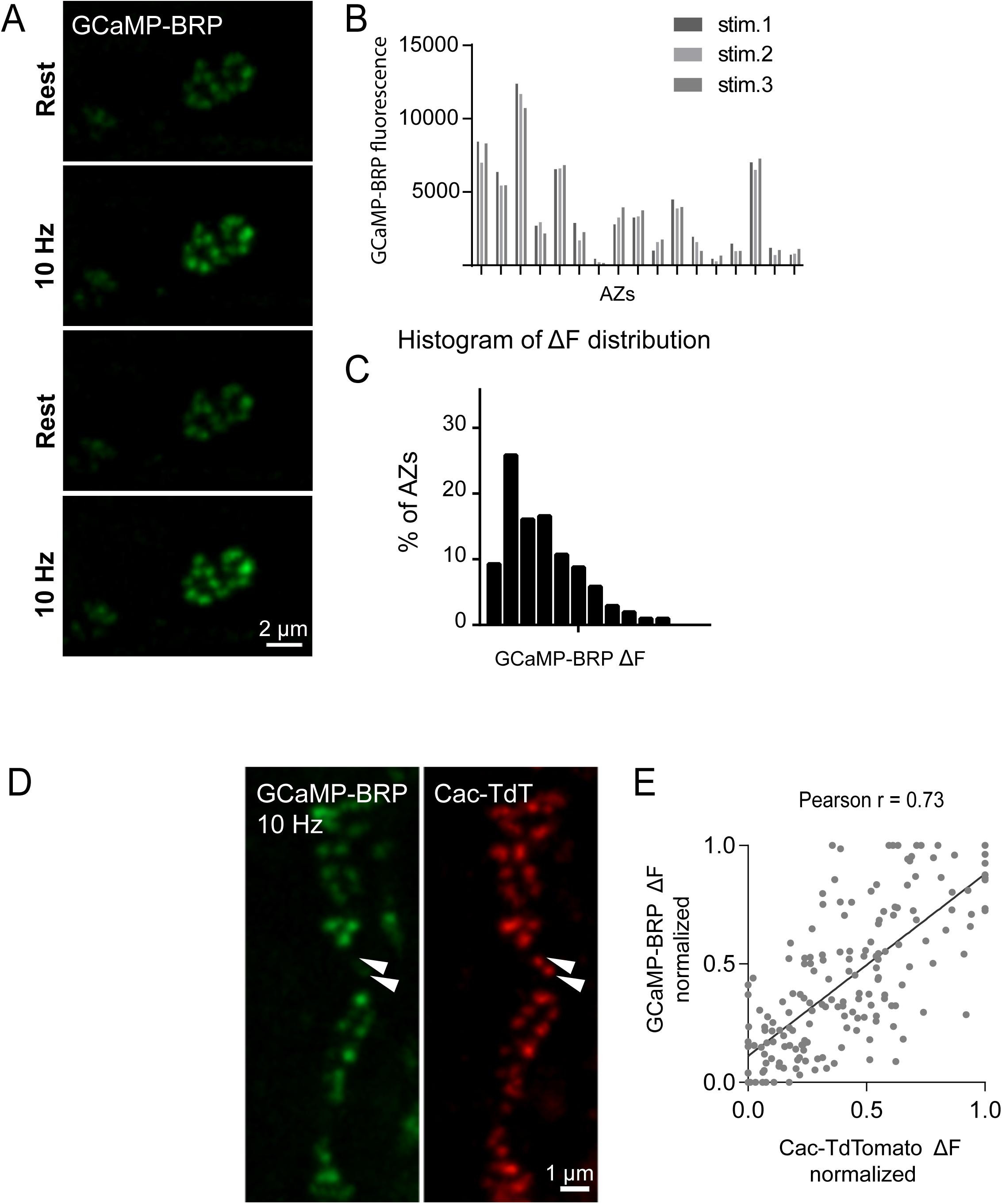
GCaMP-BRP detects relative Ca^2+^ influx at single AZs and is correlated with Cac channel density. (**A**) Representative images of the same muscle 4 NMJ bouton showing CCaMP6m-BRP fluorescence at rest and following 10 Hz stimulation for two consecutive rounds. (**B**) The AZ fluorescence intensity was plotted for three independent rounds of stimulation for BRP-GCaMP6m. Fluorescence changes per AZ remain stable for the same AZ during multiple rounds of stimulation. (**C**) Histogram of the distribution of relative fluorescence intensities (ΔF) across AZs for BRP-GCaMP6m. (**D**) Representative images showing GCaMP6m-BRP fluorescence before (left panel) and during stimulation (middle panel). The corresponding distribution of Cac channels labeled by Cac-TdTomato is shown for the same NMJ (right panel). Examples of rare Cac-positive AZs that showed no corresponding Ca^2+^ influx are indicated (arrows). (**E**) Correlation between GCaMP6m-BRP ΔF during stimulation and Cac-TdTomato fluorescence intensity at individual AZs.

To assay the ability of the sensor to detect local Ca^2+^ influx, the stability of ^GCaMP^BRP-visualized Ca^2+^ signals during multiple rounds of 5-second 10 Hz stimulation was determined. An image Z-stack was collected from individual boutons, and AZ fluorescence intensity was analyzed for each round of stimulation. Although the amount of fluorescence increase (ΔF) was different for each AZ, it was very stable at the same AZ for each independent stimulation (Figure 5B). AZs that had lower levels of Ca^2+^ influx in the first round of stimulation had similarly low ΔF across all three rounds (Figure 5B). Plotting the frequency distribution of all ΔF signals confirmed that ^GCaMP^BRP-short displayed a heterogeneous distribution of ΔF across AZs during stimulation (n = 205 AZs from 6 NMJs from 3 animals, Figure 5C).

We next assayed if Ca^2+^ influx detected by ^GCaMP^BRP-short is correlated with Cac channel density estimated by the fluorescence of Cac-TdTomato. Animals expressing both transgenes in the presynaptic compartment displayed a strong correlation (Pearson r = 0.73, R^2^ = 0.53, p<0.0001, n = 176 AZs from 7 NMJs from 6 animals) between the Ca^2+^ dependent excitation of ^GCaMP^BRP-short (ΔF) and Cac-TdTomato intensity at individual AZs during stimulation (Figure 5D, E). In contrast, a weaker correlation (Pearson r = 0.18, R^2^ = 0.03, p<0.001, n = 338 AZs from 8 NMJs from 6 animals) of ^GCaMP^BRP-short ΔF and Cac intensity at rest was observed. These data indicate that the overall density of Cac channels detected by fluorescent tagging provides a reasonable estimation of the expected Ca^2+^ influx for each AZ. However, there were some instances where specific AZs experienced a disproportionally low ΔF of ^GCaMP^BRP-short signal relative to their Cac-TdTomato intensity (Figure 5D, arrows). This observation suggests that Ca^2+^ influx can be fine-tuned and regulated independently of Ca^2+^ channel abundance at certain AZs. As such, measuring both Ca^2+^ channel density and Ca^2+^ influx is likely to provide a more accurate readout of how *P_r_* is controlled by local Ca^2+^ concentrations near the mouth of Ca^2+^ channel clusters.

Using ^GCaMP^BRP-short as a tool to estimate Ca^2+^ influx at individual AZs, we analyzed the correlation between ^GCaMP^BRP-short ΔF induced by 10 Hz stimulation and release rate visualized by myr-jRGECO1a during 1 Hz stimulation at single release sites (Figure 6A, B). AZ heatmaps for both *P_r_* and Ca^2+^ influx fluorescence intensity were generated and compared across the AZ population (Figure 6A). AZs that experienced stronger Ca^2+^ influx displayed the highest *P_r_* during stimulation. Overall, there was a strong correlation between Ca^2+^ influx and AZ *P_r_* (Pearson r = 0.56, R^2^ = 0.31, p<0.0001, n = 492 AZs from 6 NMJs from 6 animals, Figure 6B), indicating the levels of Ca^2+^ influx play a major role in determining whether a synaptic vesicle undergoes fusion during an evoked response. In contrast, the frequency of spontaneous vesicle fusion per AZ was only mildly correlated with the amount of Ca^2+^ influx detected by ^GCaMP^BRP-short (Pearson r = 0.23, R^2^ = 0.07, n = 492 AZs from 6 NMJs from 6 animals, representative experiment shown in Figure 6C), consistent with spontaneous release being largely independent of extracellular Ca^2+^ at this synapse. It is worth noting that although a strong correlation between Ca^2+^ influx and evoked *P_r_* was observed at most AZs, a minority population of AZs that displayed robust Ca^2+^ influx had very low *P_r_* (Figure 6B). In summary, these data indicate that Ca^2+^ channel density and Ca^2+^ influx are key factors regulating evoked release at individual AZs. In addition, other factors can negatively influence *P_r_* at a small minority of AZs independent of Ca^2+^ influx.

**Figure 6.**
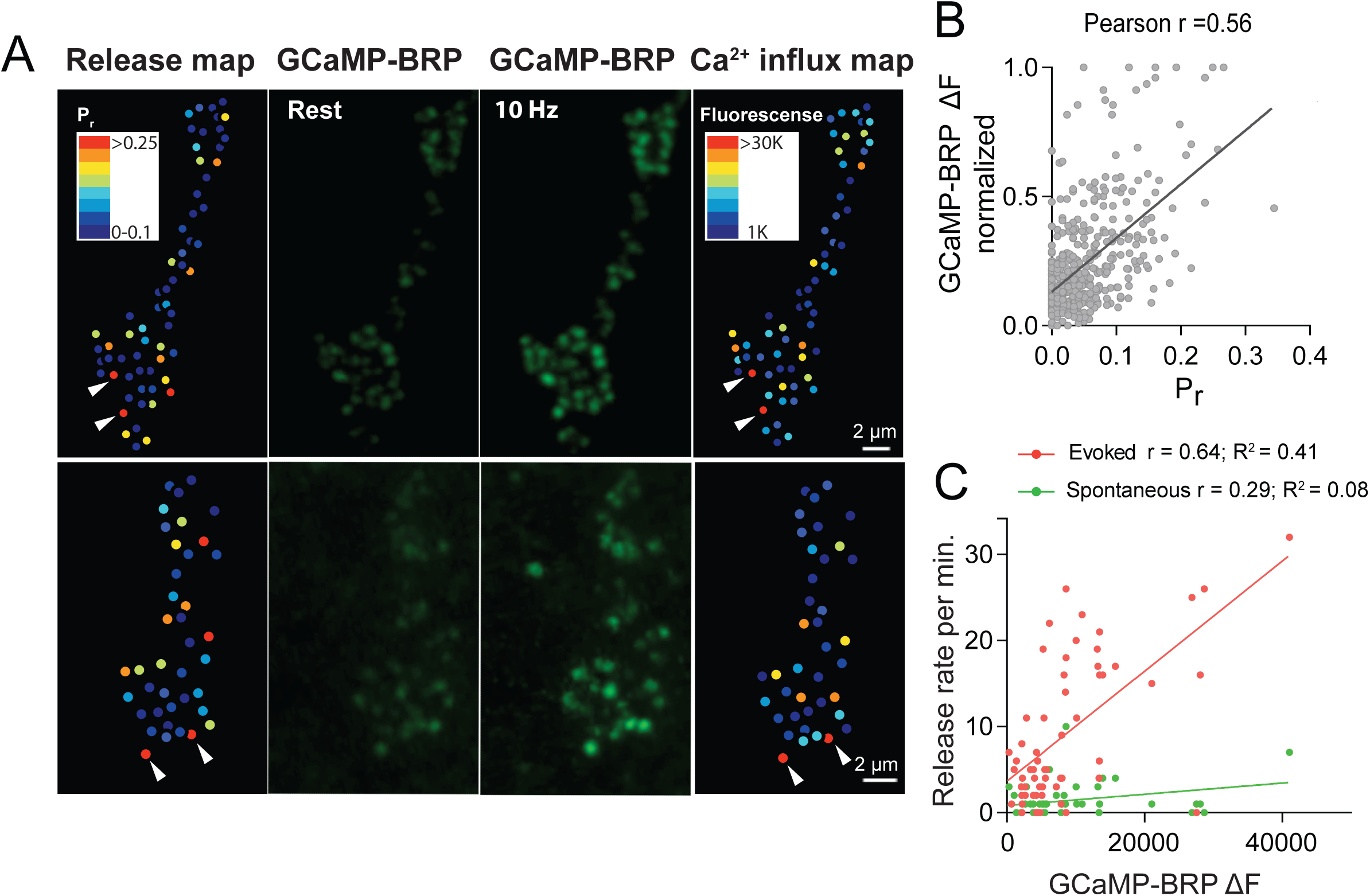
*P_r_* correlates with the relative levels of Ca^2+^ influx at AZs. (**A**) Two representative muscle 4 NMJs with AZ *P_r_* heatmaps obtained following myr-jRGECO mapping during stimulation (left panel). GCaMP6m-BRP fluorescence levels of the same NMJ at rest (second panel) and during stimulation (third panel) are shown. Heatmaps of GCaMP6m-BRP ΔF during stimulation are displayed in the right panel. Several representative high *P_r_* AZs that experienced the strongest Ca^2+^ influx detected by GCaMP-BRP are noted (arrows). (**B**) Correlation between GCaMP6m-BRP ΔF (during 10 Hz stimulation) and AZ *P_r_* (during 1 Hz stimulation) is shown across all experiments. (**C**) Representative correlation between GCaMP6m-BRP ΔF and AZ release rate per minute for evoked (red) and spontaneous (green) fusion for a representative single NMJ.

### Segregation of postsynaptic glutamate receptor subunits at high *P_r_* AZs

We next examined if glutamate receptor composition in the postsynaptic compartment varied at low *P_r_* versus high *P_r_* AZs. At the *Drosophila* NMJ, glutamate receptors assemble as heteromeric tetramers containing three essential subunits (GluRIII, IID and IIE) and a variable 4^th^ subunit of GluRIIA or GluRIIB (Featherstone et al., 2005; Marrus et al., 2004; Petersen et al., 1997; Qin et al., 2005; Schuster et al., 1991). However, it is unclear if GluRIIA and GluRIIB differentially accumulate at AZs in a manner that correlates with presynaptic *P_r_*. To visualize GluRIIA and GluRIIB, GluRIIA-RFP and GluRIIB-GFP tagged proteins were expressed under the control of their endogenous promoters. To image myrGCaMP6s activity without obscuring GluRIIB-GFP, myrGCaMP6s was expressed at low levels using the LexA/LexOP system. We generated LexAop-myrGCaMP6s transgenic animals and expressed myrGCaMP6s in muscle 4 with Mef2-LexA. LexA driven myrGCaMP6s signal is much dimmer than UAS-myrGCaMP6s. The fluorescent signal is observed at very low uniform levels in the muscle membrane in the absence of Ca^2+^ influx, and does not obscure the much brighter GluRIIB-GFP PSD puncta (Figure 7B, middle panel). However, upon Ca^2+^ binding to myrGCaMP6s, the fluorescence dramatically increases at AZs compared to the level of the endogenous GluRIIB-GFP signal, allowing simultaneous imaging of baseline GluRIIB-GFP levels and synaptic activity detected by myrGCaMP6s during stimulation.

**Figure 7.**
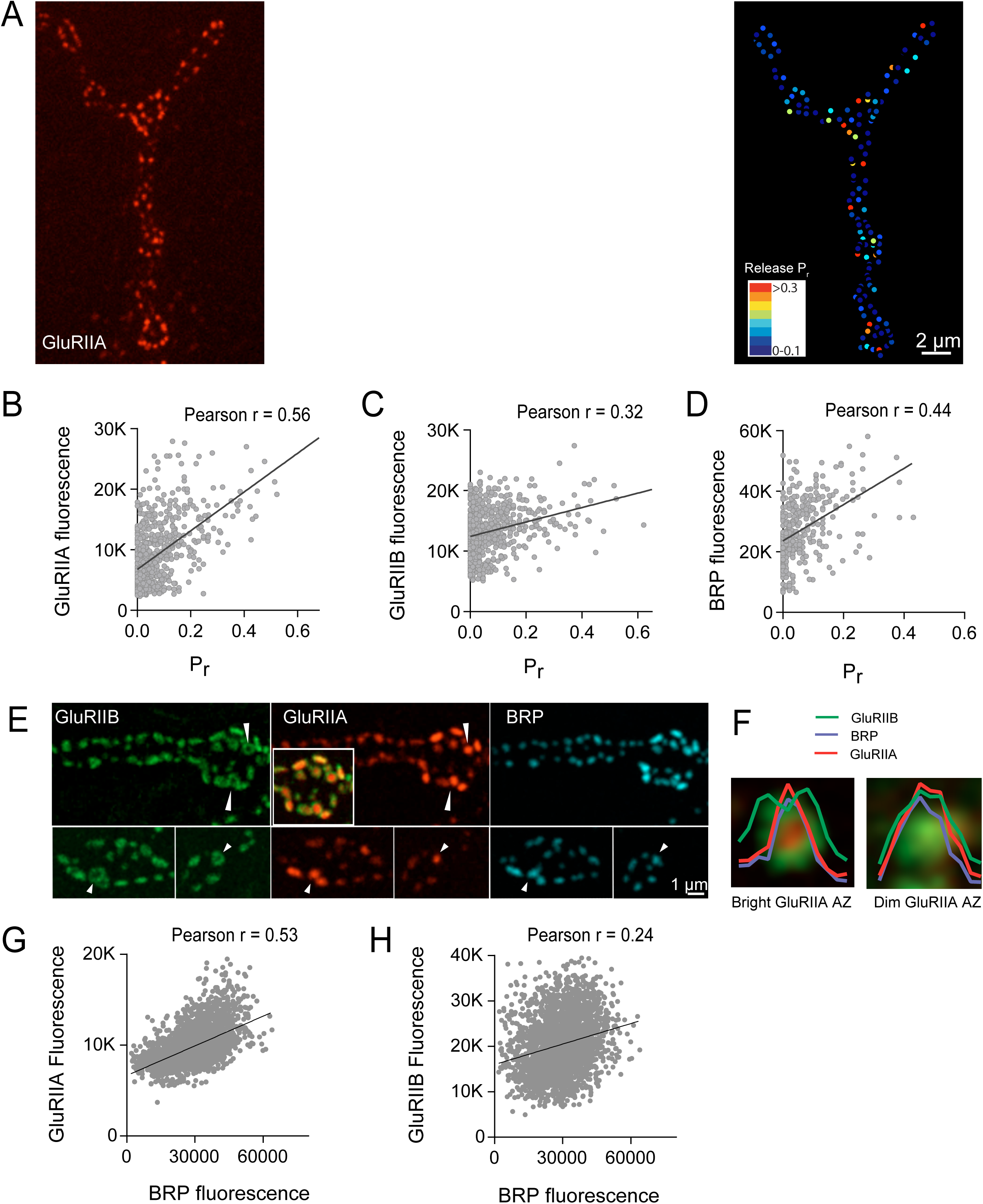
High *P_r_* AZs have elevated PSD GluRIIA levels and display a distinct pattern of glutamate receptor clustering. (**A**) Representative image showing the heterogeneous distribution of GluRIIA-RFP (left panel) at a 3^rd^ instar muscle 4 NMJ. More uniform GluRIIB-GFP PSD puncta can also be observed over the much dimmer myrGCaMP6s (second panel). BRP distribution (third panel) and *P_r_* heatmaps (right panel) for the same NMJ are shown. The correlation between AZ *P_r_* and GluRIIA-RFP (**B**), GluRIIB (**C**) and BRP (**D**) fluorescence intensity is plotted. (**E**) Representative images showing distribution of GluRIIA, GluRIIB and BRP, without co-expression of myrGCaMP6s. Synapses containing bright GluRIIA puncta have GluRIIB predominantly localized to the periphery of the PSD (arrows), surrounding a GluRIIA core. These AZs have higher BRP intensities as well. (**F**) Fluorescence line profiles showing GluRIIA, GluRIIB and BRP normalized fluorescence distribution across individual AZs. All AZs were separated into two groups according to their GluRIIA brightness, with “bright” PSDs based on their GluRIIA intensity (2 standard deviations above average). The peripheral distribution of GluRIIB around central GluRIIA cores was only obvious for bright GluRIIA-positive PSDs that were shown to be more active during stimulation. Correlation between GluRIIA-RFP (**G**) or GluRIIB-GFP (**H**) with BRP intensity at individual AZs.

Simultaneous expression of GluRIIA-RFP and GluRIIB-GFP revealed a heterogeneous distribution of each subunit across the AZ population, with GluRIIA levels far more variable than GluRIIB (Figure 7A). Indeed, similar to the relatively sparse localization of high *P_r_* AZs across the NMJ (Figure 2), a similar sparse distribution of AZs apposed by very bright GluRIIA PSDs was observed (Figure 7A). To determine if AZs that preferentially accumulate high levels of GluRIIA correspond to high *P_r_* release sites, we mapped *P_r_* across the AZ population in GluRIIA-RFP/GluRIIB-GFP expressing animals. Analysis of the *P_r_* map revealed a strong positive correlation between GluRIIA-RFP and *P_r_* (Pearson r = 0.56, R^2^ = 0.32, p<0.0001, n = 756 AZs from 8 NMJs from 4 animals, Figure 7B). In contrast, correlation with the levels of GluRIIB-GFP was weaker (Pearson r = 0.32, R^2^ = 0.1, p<0.0001, n = 756 AZs from 8 NMJs from 4 animals, Figure 7C). A correlation between brighter GluRIIA PSD puncta and higher *P_r_* sites was also observed in the analysis of *syt1* mutants (Figure 3A, arrows). These findings are consistent with previous observations that glutamate receptors preferentially cluster at AZs with high *P_r_* based on electrophysiological and morphological studies of a *Drosophila* GluRIII hypomorphic mutant (Marrus and Diantonio, 2004). We next examined how the resident AZ protein BRP differentially accumulated at AZs in the dual labeled glutamate receptor subunit lines. After mapping *P_r_* in the larvae, preparations were fixed and stained with anti-BRP antisera. As previously observed in animals lacking tagged glutamate receptors (Figure 1), a positive correlation between AZ *P_r_* and BRP levels was observed (Pearson r = 0.44, R^2^ = 0.2, p<0.0001, n = 399 AZs from 6 NMJs from 4 animals, Figure 7D). In summary, these data indicate that GluRIIA more strongly accumulates at PSDs apposing high *P_r_* AZs.

Beyond the preferential GluRIIA accumulation at high *P_r_* sites, we also observed a change in GluRIIA/GluRIIB distribution within single PSDs. The PSDs apposing the highest *P_r_* AZs showed a skewed distribution of the receptor subtypes, with GluRIIA concentrating in the center of the receptor field immediately apposing the presynaptic BRP cluster (Figure 7E, arrows). At these sites, GluRIIB occupied a more peripheral position around the central GluRIIA cluster. A similar localization pattern with a ring of GluRIIB surrounding a central GluRIIA patch was previously noted with antibody staining for the two receptors at some larval AZs in late 3^rd^ larvae (Marrus et al., 2004). To analyze this segregation in glutamate receptor distribution at single PSDs in greater detail, GluRIIA/B staining was examined in the absence of co-expressed myrGCaMP6s to avoid any overlap in the GFP channel. Our prior analysis (Fig. 7B) indicated the brightest GluRIIA PSDs correspond to high *P_r_* sites. Bright PSDs were selected based on their GluRIIA intensity (2 standard deviations above average) and line profiles were drawn across each PSD defined by their corresponding presynaptic BRP puncta. The intensity of pixels along that line for each fluorophore was then analyzed. Average pixel intensity revealed drastically distinct profiles for GluRIIB distribution between “bright” and “dim” PSDs classified based on their GluRIIA intensity. GluRIIB was more evenly distributed across the entire PSD at dim GluRIIA sites, but was segregated outward, forming a circular donut-like ring around central GluRIIA puncta at bright GluRIIA sites (Figure 7F). In addition, presynaptic BRP intensity was more strongly correlated with postsynaptic GluRIIA levels (Pearson r = 0.53, R^2^ = 0.28, p<0.0001, n = 2496 AZs from 19 NMJs from 7 animals, Figure 7G) compared to GluRIIB (Pearson r = 0.24, R^2^ = 0.05, p<0.0001, n = 2496 AZs from 19 NMJs from 7 animals, Figure 7H). Overall, these findings indicate the postsynaptic cell accumulates GluRIIA and redistributes GluRIIB to the PSD periphery at high *P_r_* sites.

### Analysis of *Pr* acquisition during AZ development using glutamate receptor segregation as a proxy

The *Drosophila* larval NMJ is a highly dynamic structure, with new synaptic boutons and AZs undergoing continuous addition throughout development (Harris and Littleton, 2015; Rasse et al., 2005; Schuster et al., 1996; Zito et al., 1999). Given the correlation between Ca^2+^ channel density, GluRIIA/GluRIIB segregation and high *P_r_*, we were interested in determining how AZs acquire a specific *P_r_* during a larval developmental period that lasts 6–7 days. One model is that certain AZs gain a higher *P_r_* status during development through a tagging or activity-dependent mechanism that would lead to preferential accumulation of key AZ components compared to their neighbors. Alternatively, high *P_r_* AZs might simply be more mature than their low *P_r_* neighbors, having an earlier birthdate and a longer timeframe to accumulate AZ material. To differentiate between these models for release heterogeneity, it would be desirable to follow *P_r_* development from the embryonic through larval stages. However, this is not technically feasible due to the small size of AZs and the rapid locomotion that larvae undergo, preventing generation of *P_r_* maps in intact moving animals. Instead, we employed an alternative approach to repeatedly image the same NMJ at muscle 26 directly through the cuticle of intact larvae during anesthesia (Andlauer and Sigrist, 2012; Fouquet et al., 2009; Füger et al., 2007; Rasse et al., 2005; Zhang et al., 2010). Using this technique, we found that anesthesia eliminated action potential induced release and the associated GCaMP signals, preventing direct *P_r_* measurements in anesthetized larvae. We instead focused on imaging GluRIIA accumulation and GluRIIA/GluRIIB segregation, which was strongly correlated with *P_r_* (Figure 7), as a proxy for the emergence of high *P_r_* sites. Prior studies demonstrate GluRIIA at PSDs closely tracks with Cac accumulation at corresponding AZs (Fouquet et al., 2009; Rasse et al., 2005), indicating the two compartments are likely to mature at similar rates. To examine this directly, we assayed whether GluRIIA and Cac accumulation were correlated during development. Indeed, the intensity of Cac-GFP and GluRIIA-RFP puncta were strongly correlated at individual AZs during early larval development (Pearson r = 0.823, R^2^ = 0.6771, p<0.0001, n= 441 AZs from 8 NMJs from 8 larvae, Figure 7 – figure supplement 1A, B), indicating GluRIIA provides a robust marker that reflects the corresponding levels of presynaptic Cac at individual AZs.

Previously described *in vivo* imaging approaches with anesthesia at *Drosophila* NMJs employed early 3^rd^ instar larvae as the starting time point, and followed the distribution of fluorescently-labeled synaptic proteins during the final ~36 hours of development prior to pupation (Rasse et al., 2005). To follow AZ *P_r_* development soon after synapse formation, we modified these techniques to allow imaging of glutamate receptor distribution at earlier stages of development (see methods). This allowed successful birth dating and successive imaging of the same AZ over a 6-day period beginning shortly after synapse formation in the early 1^st^ instar period through the late 3^rd^ instar stage (Figure 8, Figure 8 – figure supplement 1, Figure 8 – figure supplement 2). In early 1^st^ instar larvae (within 12 hours of hatching) GluRIIA and GluRIIB were largely co-localized at postsynaptic puncta (Figure 8, Figure 8 – figure supplement 1). One exception was the presence of diffuse GluRIIA that accumulated around unusually long axonal extensions that emerged from presynaptic boutons (Figure 8A, arrows). These structures were devoid of any detectable GluRIIB or the bright GluRIIA puncta that are associated with AZs, and may be remnants of previously described muscle filopodial structures, termed myopodia, that interact with presynaptic filopodia to dynamically regulate early synaptic target recognition (Kohsaka and Nose, 2009; Ritzenthaler et al., 2000; Ritzenthaler and Chiba, 2003). GluRIIA appears concentrated on these structures, as has been observed for the leucine-rich repeat cell adhesion protein Capricious (Kohsaka and Nose, 2009). Repeated imaging of these thinner GluRIIA-positive processes revealed that they were capable of developing into mature synaptic boutons with concentrated GluRIIA and GluRIIB synaptic puncta (Figure 8 – figure supplement 3). By 24 hours of larval growth, GluRIIA rich extensions were no longer observed, indicating these structures are restricted to early developmental stages.

**Figure 8.**
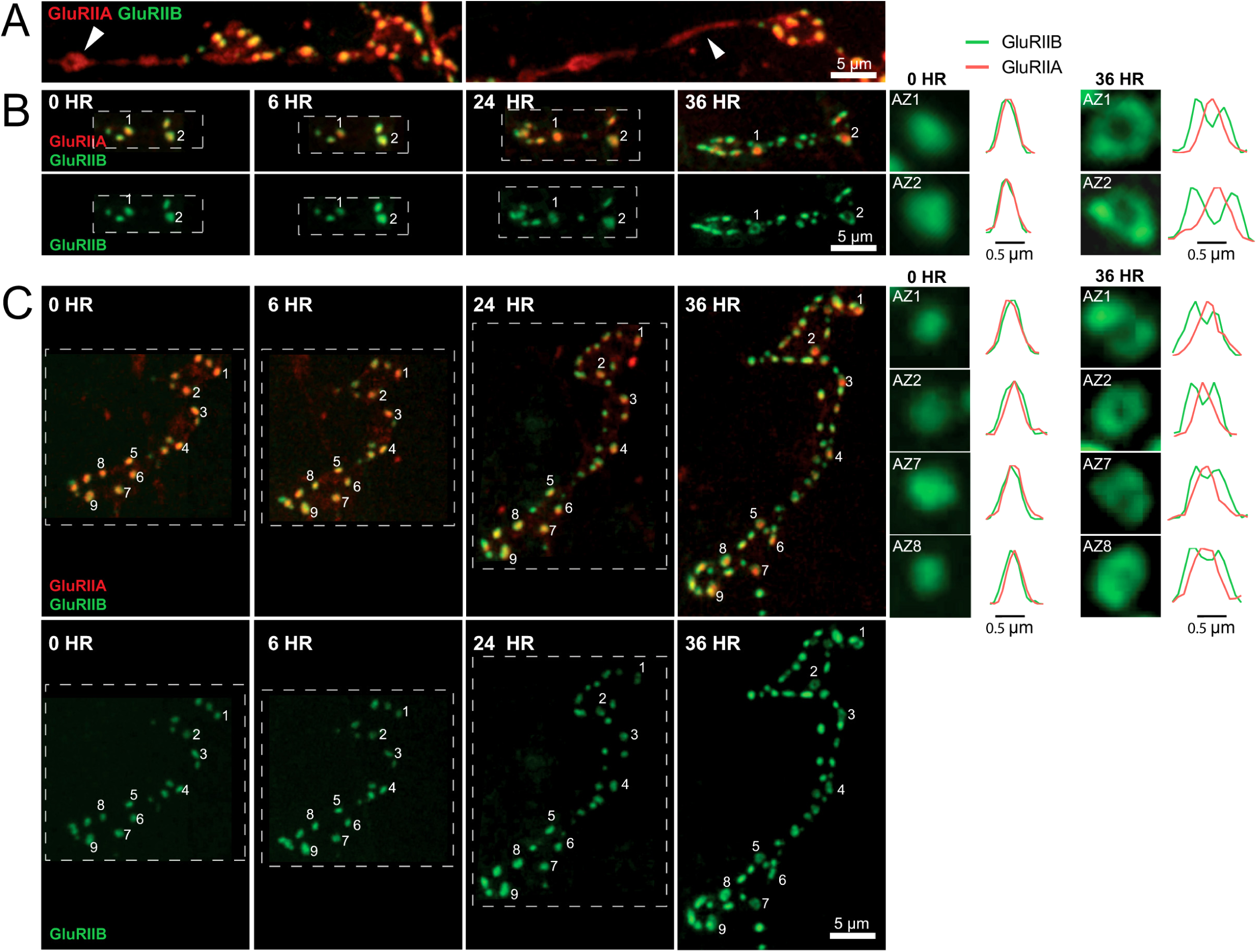
Glutamate receptor segregation during PSD development. **(A**) Two representative muscle 26 NMJs visualized during live imaging in early 1^st^ instar larvae expressing GluRIIA-RFP and GluRIIB-GFP. The arrows denote GluRIIA-RFP positive extensions from the main arbor that are devoid of detectable PSDs or GluRIIB at this stage of development. These extensions disappear during imaging from later larval stages, but some go on to develop fully formed synaptic boutons with new AZs (Figure 8 – supplemental figure 3). (**B**) Representative serial time points of NMJ development visualized by repeated imaging through the cuticle of an anesthetized larvae at the indicated time points beginning at the early 1^st^ instar stage. Two of the five PSDs present during the first imaging session are labeled and are the first to develop the peripheral GluRIIB segregation pattern 36 hours later. GluRIIB labeling alone is shown in the bottom panel. The right panels show GluRIIB fluorescence and normalized GluRIIA and GluRIIB fluorescent line profiles for the indicated PSDs at the initial imaging session (0 hour) and 36 hours later. (**C**) Serial images of an NMJ with a larger number of AZs present at the 1^st^ instar stage. After 36 hours of development, the peripheral segregation of GluRIIB around GluRIIA was first observed at PSDs that were present during the initial imaging session (numbered). The right panels show GluRIIB fluorescence and normalized GluRIIA and GluRIIB fluorescent line profiles for the indicated PSDs at the initial imaging session (0 hour) and 36 hours later. The dashed box surrounds the actual imaged segment of the NMJ in each panel.

Live imaging of GluRIIA and GluRIIB distribution at early PSDs in anesthetized 1^st^ instar larvae demonstrated that the receptors were co-localized and lacked the segregation where GluRIIB clustered around central GluRIIA puncta that was observed at high *P_r_* sites later in development (Figure 8B). The first emergence of GluRIIA/B segregation, with GluRIIB rings surrounding a GluRIIA core, was observed after 36 hours of imaging from the 1^st^ instar period (Figure 8B, C). The GluRIIA/B segregation always emerged first at the oldest and most mature synapses that existed previously on the 1^st^ day of imaging (Figure 8B, C). The most mature PSDs also contained more GluRIIA fluorescent signal (17430 ± 634.0, n = 86 AZs from 8 NMJs from 5 animals) compared to later born synapses that emerged during the 48 hour imaging session (8909 ± 289.8, n = 210 AZs from 8 NMJs from 5 animals). During later larval development, the cuticle thickness changed dramatically and prevented reliable comparison of absolute receptor density with earlier stages. However it was clear that GluRIIA intensities that were uniform in 1^st^ instar larvae became more heterogeneous at the 3^rd^ instar stage (Figure 8 – figure supplement 2). Indeed, histograms of normalized fluorescence intensity (relative intensity scaled from 0 to 1) revealed that GluRIIA and GluRIIB were distributed relatively uniformly at 1^st^ instar larval PSDs, with GluRIIA distribution becoming more skewed at later stages (Figure 9A, B). These results indicate that GluRIIA/GluRIIB fluorescence distribution is highly heterogeneous by the early 3^rd^ instar stage, with the brightest GluRIIA positive PSDs, and by extension their corresponding high *P_r_* AZs, representing those that appeared earliest in development.

**Figure 9.**
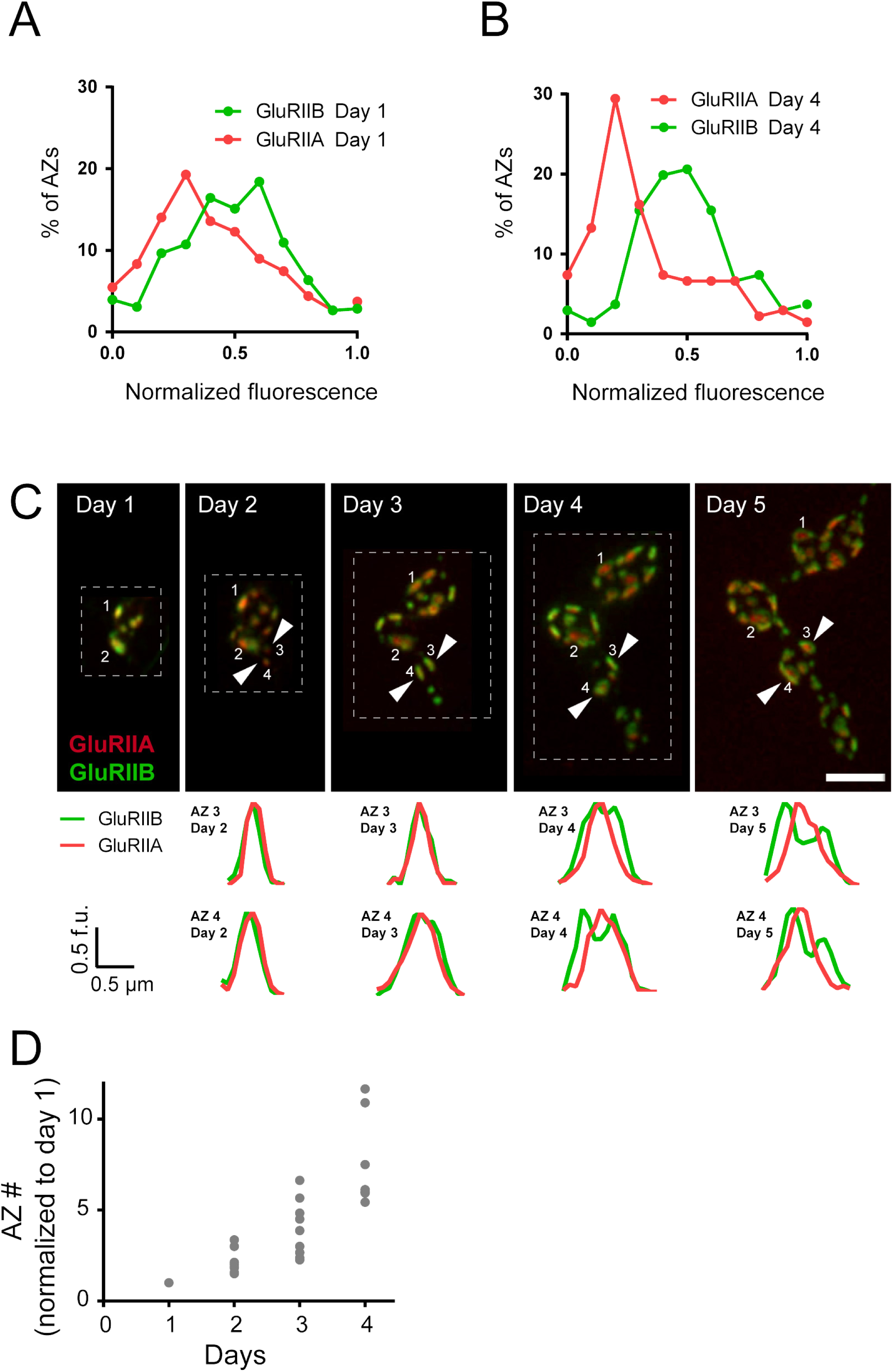
Rate of acquisition of glutamate receptor segregation during development. Histograms of the distribution of normalized GluRIIA and GluRIIB fluorescence at the 1^st^ instar (day 1) (**A**) and 3^rd^ instar (day 4) (**B**) stages for muscle 26 imaged through the cuticle of anesthetized larvae. For each data set, GluRIIA and GluRIIB fluorescence is presented from dimmest (0) to brightest (1). GluRIIA shows a more skewed distribution of fluorescence at day 4, consistent with its accumulation at high *P_r_* AZs. (**C**) Representative muscle 26 NMJ image sequence showing appearance and maturation of two new synapses (#3 and 4) that were not present in the initial imaging session. Several preexisting synapses (#1 and 2) that developed the typical GluRIIB donut structure later in development are also labeled. The dashed box surrounds the actual imaged segment of the NMJ. New GluRIIA and GluRIIB clusters appear initially as small puncta (day 2, arrows) that become brighter on day 3. By day 4 they begin to display the donut like GluRIIB profile. At day 5, GluRIIB distribution to the periphery around a bright GluRIIA PSD representative of high *P_r_* sites becomes prominent. The bottom panels show normalized GluRIIA and GluRIIB fluorescent line profiles for the newly identified PSDs (#3, 4) throughout the 5 day imaging series. (**D**) Changes in AZ number during larval maturation at muscle 26 presented as a ratio of AZs observed during the first day of imaging (day 1).

Over what time frame do synapses developmentally acquire high *P_r_* properties? To estimate the average time required for this process, the interval from the first emergence of a synapse in an imaging session to the time point when segregation of GluRIIB around GluRIIA central puncta occurred was calculated. This analysis was restricted to newly formed AZs that appeared during the imaging sessions, and excluded AZs that were present in the first imaging session performed in 1^st^ instar larvae. The average time from the first emergence of a synapse to when it acquired the segregated GluRIIA/B pattern observed at high *P_r_* AZs was 3.20 ± 0.08 days (n = 41 AZs from 7 NMJs from 3 animals, Figure 9C). In a small subset of PSDs (5%), a slightly faster accumulation of GluRIIA and the formation of GluRIIB peripheral rings was observed, but never faster than 2 days (Figure 8 – figure supplemental 2). Given that the developing NMJ is adding AZs at a rapid rate (Rasse et al., 2005; Schuster et al., 1996), we estimated if AZ maturation time identified over the course of our live imaging experiments would fit with the 9.9% of high *P_r_* sites observed at the early 3^rd^ instar stage by GCaMP imaging. The number of synapses present at the same NMJ from the 1^st^ instar through the early 3^rd^ instar stage was quantified from live imaging experiments (Figure 9D). AZ number doubled each day, such that the average number of AZs found at the 1^st^ instar stage (day 1) represented 14.7 ± 1.4% (n = 8 NMJs from 3 animals) of all AZs present by day 4 (3 days after initial imaging in 1^st^ instars). This value is similar to the 10% population of high *P_r_* AZs observed during *P_r_* mapping. Overall, these data support the hypothesis that AZ maturation is a key factor in regulating *P_r_*, leading to increased accumulation of Ca^2+^ channels and GluRIIA/GluRIIB segregation at high *P_r_* sites compared to AZs that are newly formed (<2 days).

## Discussion

In the current study we used optical quantal imaging to examine the source of heterogeneity in evoked *P_r_* across the AZ population at *Drosophila* NMJs. By combining quantal imaging with SIM microscopy, we first confirmed that release heterogeneity was not caused by summation of fusion events from multiple AZs. By monitoring release over 15 minute intervals, we also observed that *Pr* was a stable feature of each AZ. The *Drosophila* genome encodes a single member of the N/P/Q-type Ca^2+^ channel a1 subunit family (Cac) that is present at AZs and is responsible for neurotransmitter release (Fouquet et al., 2009; Kawasaki et al., 2004, 2000; Littleton and Ganetzky, 2000; Liu et al., 2011; Rieckhof et al., 2003; Smith et al., 1996). Using transgenically labeled Cac lines, we found that Cac density at AZs strongly correlated with *P_r_*. To directly visualize presynaptic Ca^2+^ influx at single AZs, GCaMP fusions to the core AZ component BRP (the *Drosophila* ELKS/CAST homolog) were generated. The levels of Ca^2+^ influx at single AZs was highly correlated with both Cac density and *P_r_*. In contrast, loss of the Synaptotagmin 1 synaptic vesicle Ca^2+^ sensor did not change the heterogeneous distribution of *P_r_* across AZs.

High *P_r_* AZs also displayed a distinct pattern of glutamate receptor clustering. Most PSDs showed a homogeneous distribution of the GluRIIA and GluRIIB containing subunits on the apposing postsynaptic membrane. In contrast, high *P_r_* AZs were apposed by PSDs where GluRIIA receptors concentrated at the center of the AZ, with GluRIIB receptors forming a ring at the PSD periphery. A similar activity-dependent segregation of GluRIIA and a GluRIIA gating mutant has been previously observed at individual AZs in *Drosophila* (Petzoldt et al., 2014). In addition, anti-glutamate receptor antibody staining of wildtype larvae lacking tagged glutamate receptors identified a GluRIIB ring around the GluRIIA core in some mature 3^rd^ instar NMJ AZs (Marrus et al., 2004). The correlation of *P_r_* with GluRIIA accumulation is especially intriguing considering that this subunit has been implicated in homeostatic and activity-dependent plasticity (Davis, 2006; Frank, 2014; Petersen et al., 1997; Sigrist et al., 2003). By following the developmental acquisition of this postsynaptic property as a proxy for *P_r_* from the 1^st^ through 3^rd^ instar larval stages via live imaging, we observed that the earliest formed AZs are the first to acquire this high *P_r_* signature over a time course of ~3 days.

Similar to our prior observations (Melom et al., 2013), we found that most AZs at the *Drosophila* NMJ have a low *P_r_*. For the current study, the AZ pool was artificially segregated into low and high release sites, with high releasing sites defined based on having a release rate greater than two standard deviations above the mean. Given that birthdate is a key predictor of glutamate receptor segregation, and by proxy *P_r_*, we expect the AZ pool to actually reflect a continuum of *P_r_* values based on their developmental history. However, using the two standard deviation criteria, 9.9% of AZs fell into the high *P_r_* category, with an average *P_r_* of 0.28 in extracellular saline containing 1.3 mM Ca^2+^ and 20 mM MgCl_2_. The remaining AZs that participated in evoked release had an average *P_r_* of 0.05. Ca^2+^ channel density and Ca^2+^ influx at individual AZs was a key determinant of *P_r_* heterogeneity, as evoked *P_r_* and the density of Cac channels tagged with either TdTomato or GFP displayed a strong positive correlation (Pearson r = 0.65 and Pearson r = 0.58, respectively). To bypass any unknown effects of channel tagging on *P_r_*, we also created a tool to directly visualize AZ Ca^2+^ influx by tethering GCaMP to the AZ protein BRP. Again, a strong correlation between Ca^2+^ influx and AZ *P_r_* (Pearson r = 0.60) was observed, confirming that the levels of Ca^2+^ influx are a major determinant for synaptic vesicle fusion during an evoked response. Spontaneous fusion showed a much weaker correlation with both Cac density and Ca^2+^ influx at individual AZs, consistent with prior studies indicating spontaneous release rates are poorly correlated with external Ca^2+^ levels at this synapse (Jorquera et al., 2012; Lee et al., 2013). With synaptic vesicle fusion showing a steep non-linear dependence upon external Ca^2+^ with a slope of ~3–4 (Dodge and Rahamimoff, 1967; Heidelberger et al., 1994; Jan and Jan, 1976), a robust change in *P_r_* could occur secondary to a relatively modest increase in Ca^2+^ channel density over development. Although the number of Ca^2+^ channels at a *Drosophila* NMJ AZ is unknown, estimates of Cac-GFP fluorescence during quantal imaging indicate a ~2-fold increase in channel number would be necessary to move a low *P_r_* AZ into the high *P_r_* category. Similar correlations between evoked *P_r_* and Ca^2+^ channel density have been found at mammalian synapses (Holderith et al., 2012; Nakamura et al., 2015; Sheng et al., 2012), suggesting this may represent a common mechanism for determining release strength at synapses.

Beyond low and high *P_r_* sites, we found that 9.7% of the AZs analyzed displayed only spontaneous release. We could detect no fusion events for either evoked or spontaneous release for another 14.6% of AZs that were defined by a GluRIIA-positive PSD in live imaging. Whether these cases represent immature AZs with extremely low evoked *P_r_*, or distinct categories reflective of differences in AZ content, is unknown. For spontaneous-only sites, we previously found that the ΔF/F_avg_ quantal signal detected postsynaptically by GCaMP imaging was similar to that observed at mixed mode AZs displaying both evoked and spontaneous events, indicating that there is unlikely to be a dramatic difference in glutamate receptor density at these sites versus low *P_r_* AZs (Melom et al., 2013). Further categorization of spontaneous-only AZs and silent AZs will require the identification of unique makers that can be used to track them in live imaging during synaptic development.

Other factors we examined for regulating *Pr* were differences in local synaptic vesicle pools or synaptic vesicle protein content or state (for example, phosphorylation). *Pr* was largely unchanged with either 5-minute rest or 5 minute 5 Hz stimulation between imaging sessions. Although *Pr* may be more dynamic over longer intervals, the observation that developmental maturation of glutamate receptor segregation occurs over ~ 3 days is consistent with *P_r_* being a stable feature of the AZ over shorter time periods (hours to ~1 day). To further examine the role of potential heterogeneity due to differences in synaptic vesicle proteins, we assayed whether *P_r_* heterogeneity was abolished in mutants lacking the Ca^2+^ sensor Synaptotagmin 1, a major regulator of *P_r_* (DiAntonio and Schwarz, 1994; Fernández-Chacón et al., 2001; Geppert et al., 1994; Littleton et al., 1994, 1993; Yoshihara and Littleton, 2002). GCaMP imaging confirmed that release was dramatically reduced and largely asynchronous in *syt1* nulls, and that mini frequency per AZ was increased. Although *P_r_* was reduced in *syt1, P_r_* distribution among different AZs remained heterogeneous, suggesting that the AZ, rather than differential distribution of Syt1, is critical. Although roles for other synaptic vesicle proteins in *P_r_* heterogeneity cannot be excluded, the observation that prolonged 5 Hz stimulation, which would be predicted to turn over the synaptic vesicle pool, does not change *P_r_* argues against this hypothesis. Instead, these data support a model that differences in the abundance of presynaptic Ca^2+^ channels underlies heterogeneous AZ *P_r_*.

We considered several models for how AZs acquire this heterogeneous nature of *P_r_* distribution. One possibility is that unique AZs gain high *P_r_* status through a mechanism that would result in preferential accumulation of key AZ components compared to their neighbors. Given that retrograde signaling from the muscle is known to drive synaptic development at *Drosophila* NMJs (Ball et al., 2010; Berke et al., 2013; Harris and Littleton, 2015; Keshishian and Kim, 2004; McCabe et al., 2003; Piccioli and Littleton, 2014; Yoshihara et al., 2005), certain AZ populations might have preferential access to specific signaling factors that would alter their *P_r_* state. Another model is that AZs compete for key presynaptic *P_r_* regulators through an activity-dependent process. Once an AZ achieves a slight imbalance over its neighbors in release strength, it would then “win” and preferentially accumulate AZ material, similar to the concept of synapse elimination during motor neuron competition at mammalian NMJs (Tapia et al., 2012). Finally, high *P_r_* AZs might simply be more mature than their low *P_r_* neighbors, having a longer timeframe to accumulate AZ components. This model would not require any AZ to gain a favored status. AZs that appeared first during development would have more time to accumulate AZ material at a rate that would be largely identical over all AZs. Given that the *Drosophila* NMJ is constantly forming new AZs at a rapid pace during development (Rasse et al., 2005; Schuster et al., 1996), newly formed AZs would take longer to mature, generating a population of “older” AZs that would increase their *P_r_* at the same proportional rate as their newer neighbors. Given that GCaMP imaging indicates high *P_r_* sites segregate GluRIIA and GluRIIB differently from low *P_r_* sites, with the IIA isoform preferentially localizing at the center of PSDs apposing high *P_r_* AZs, we used developmental acquisition of this property as an indicator of high *P_r_* sites. Although segregation of glutamate receptors may not perfectly replicate the timing of *P_r_* acquisition during development, it is currently the best tool we have at our disposal for sequential live imaging. Based on the acquisition of GluRIIA/GluRIIB segregation, our data support the model that increases in *P_r_* reflect a time-dependent maturation process at the NMJ. Indeed, AZs that developed later in imaging sessions showed a similar time course for acquiring the postsynaptic segregation of glutamate receptors, indicating a fixed developmental maturation from low to high *P_r_* AZs that is likely to account for the majority of release heterogeneity at this connection. The continuous addition of new AZs, which double during each day of development, ensures that the overall ratio of high to low *P_r_* sites remain relatively fixed at a low percentage, depending on developmental stage and the rate of new AZ addition.

We did not test the correlation of *P_r_* with other AZ proteins besides Cac and BRP, but it would not be surprising to see a positive correlation with the density of many AZ proteins based on the observation that maturation time is a key determinant for *P_r_*. AZ maturation is also likely to promote increased synaptic vesicle docking and availability, consistent with observations that correlate active zone size with either *P_r_* or the readily releasable pool (Han et al., 2011; Holderith et al., 2012; Matkovic et al., 2013; Matz et al., 2010; Nakamura et al., 2015; Schikorski and Stevens, 1997; Wadel et al., 2007). Although it is poorly understood how AZs are assembled during development, our data would not support a model that AZs are fully preassembled during transport and then deposited as a single “quantal” entity onto the presynaptic membrane. Rather, these data support a model of seeding of AZ material that increases developmentally over time as AZs matures, consistent with several studies of AZ development in *Drosophila* (Böhme et al., 2016; Fouquet et al., 2009). In theory, each AZ would have equal access to new AZ material, and accumulate it at a fairly constant rate, with birthdate being the primary factor in how much AZ material they contain, and correspondingly, their *P_r_* status. Although no evidence for rapid changes in *P_r_* were detected in the steady-state conditions used in the current study, homeostatic plasticity is known to alter *P_r_* over a rapid time frame (~10 minutes) at the NMJ (Davis and Müller, 2015; Frank, 2014; Frank et al., 2006). It will be interesting to determine if Cac density can change over such a rapid window, or whether the enhanced release is mediated solely through changes in Cac function and Ca^2+^ influx (Müller and Davis, 2012). Changes in the temporal order of *P_r_* development could also occur secondary to altered transport or capture of AZ material. For example, the large NMJ on muscle fibers 6 and 7 displays a gradient in synaptic transmission, with terminal branch boutons often showing a larger population of higher *P_r_* AZs (Guerrero et al., 2005; Peled and Isacoff, 2011). If AZ material is not captured by earlier synapses along the arbor, it would be predicted to accumulate in terminal boutons, potentially allowing these AZs greater access to key components, and subsequently increasing their rate of *P_r_* acquisition. In summary, our data indicate that Ca^2+^ channel density and Ca^2+^ influx at single AZs is a key determinant for release heterogeneity, and that developmental AZ maturation is a key factor in *P_r_* at the *Drosophila* NMJ.

## Materials and methods

### Drosophila stocks

Flies were cultured at 25°C on standard medium. Actively crawling 3^rd^ instar male and female larvae dwelling on top of the food were used for experiments unless noted. The following strains were used: UAS-myrGCaMP6s, UAS-GCaMP6m-BRPshort, pBid-lexAop-myrGcaMP6s, UAS-myrjRGECO; Elav–GAL4, Mef2–GAL4, UAS-CacGFP (provided by Richard Ordway); UAS-CacTdTomato (provided by Richard Ordway); GluRIIA-RFP (provided by Stephan Sigrist), GluRIIB-GFP (provided by Stephan Sigrist) and 44H10-LexAp65 (provided by Gerald Rubin). *syt1* null mutants were generated by crossing *syt1^N13^*, an intragenic *syt1* deficiency (Littleton et al., 1994), with *syt1^AD4^*, which truncates Syt1 before the transmembrane domain (DiAntonio and Schwarz, 1994).

### Transgenic constructs

The fluorescent Ca^2+^ sensor GCaMP6s was tethered to the plasma membrane with an N-terminal myristoylation (myr) sequence. The first 90 amino acids of Src64b, containing a myristoylation target sequence, were subcloned into pBID-UASc with EcoRI and BglII (creating pBID-UASc-myr). GCaMP6s cDNA (Addgene plasmid 40753) was cloned into pBID-UASc-myr with BglII and XbaI. To generate the UAS-GCaMP6m-Brp-short line, GCaMP6m (Addgene plasmid 40754) cDNA and Brp-short (gift from Dr. Tobias Rasse) were PCR amplified and double digested with EcoRI/BglII and BglII/XbaI, respectively. The two cDNA fragments were ligated and the product was used to PCR amplify the fused GCaMP6m-Brp-short cDNA. The PCR product was inserted into the vector backbone pBID-UASc after digestion with EcoRI and XbaI to generate the final plasmid pBID-UASc-GCaMP6m-Brp-short. To create UAS-myrjRGECO, the vector backbone pBID-UASc-myr was digested with BglII and XbaI. jRGECO sequence was amplified from plasmid pGP-CMV-NES-jRGECO1a (gift from Dr. Douglas Kim, Addgene plasmid #61563). The digested backbone and insert were fused according to the Gibson assembly protocol using NEBuilder HighFidelity DNA Assembly Cloning Kit (E5520). To generate pBid-lexAop-myrGcaMP6s, myrGCaMP6s was amplified by PCR and inserted into pBiD-lexAop-DSCP (gift from Brian McCabe) between NotI and XbaI sites. All transgenic *Drosophila* strains were generated by BestGene.

### Immunocytochemistry

Wandering 3^rd^ instar larvae were dissected in Ca^2+^-free HL3 solution and fixed in 4% paraformaldehyde for 10 minutes, washed in PBT (PBS plus 0.1% Triton X-100) and blocked in 5% normal goat serum (NGS) and 5% BSA in PBT for 15 minutes. Samples were incubated overnight with anti-BRP (NC82, 1:200) from the Developmental Studies Hybridoma Bank, washed for 1 hour in PBS and then incubated for 2–3 hours with Alexa Fluor 607-conjugated anti-mouse IgG at 1:1000 (Invitrogen).

### Confocal imaging and data analysis

Confocal images were obtained on a Zeiss Axio Imager 2 equipped with a spinning-disk confocal head (CSU-X1; Yokagawa) and ImagEM X2 EM-CCD camera (Hammamatsu). An Olympus LUMFL N 60X objective with a 1.10 NA was used to acquire GCaMP6s imaging data at 7 to 8 Hz. A Zeiss pan-APOCHROMAT 63X objective with 1.40 NA was used for imaging stained or live animals. 3^rd^ instar larvae were dissected in Ca^2+^-free HL3 containing 20 mM MgCl_2_. After dissection, preparations were maintained in HL3 with 20 mM MgCl_2_ and 1.3 mM Ca^2+^ for 5 minutes. To stimulate the NMJ, motor nerves were cut close to the ventral ganglion and sucked into a pipette. Single pulses of current were delivered every one second for myr-jRGECO mapping or every three seconds for GCaMP6s mapping with an AMPI Master-8 stimulator using a stimulus strength just above the threshold for evoking EJPs. A 3D image stack was taken before the GCaMP imaging session to generate a full map of GluRIIA or Cac channel distribution. Later, single focal planes were imaged continuously for 4-5 minutes to collect GCaMP signals. Volocity 3D Image Analysis software (PerkinElmer) was used to analyze images. All images were Gaussian filtered (fine) to reduce noise and a movement-correction algorithm was applied. To enhance identification of myrGCaMP6 flashes, background myrGCaMP fluorescence was subtracted by creating a composite stack of 5–6 images during intervals when no synaptic release occurred. To identify the position of GluRIIA receptors and corresponding Ca^2+^ events, a 3D stack image of GluRIIA was merged to create a single plane. AZ position was identified using the “find spot” algorithm in Volocity 3.2 software that detects fluorescent peaks. ROIs with identical 5-pixel size (0.138 μm/pixel) were automatically generated by the software from identified GluRIIA spots. All GCaMP flashes were detected using the intensity threshold tool and assigned to specific ROIs based on proximity of their centroids. The time and location of Ca^2+^ events were imported into Excel or Matlab for further analysis. The number of observed GCaMP events per AZ was divided by the number of delivered stimuli to calculate AZ *P_r_*. Analysis of Cac, BRP, GluRIIA or GluRIIB intensities was performed similarly, identifying AZ fluorescence peaks and defining 3 pixel square ROIs around each peak to calculate average fluorescence. Average AZ fluorescence intensities of 3-pixel square ROIs was also used for correlation analysis.

### Live imaging

Larvae were anesthetized with SUPRANE (desflurane, USP) from Amerinet Choice (Zhang et al., 2010). Larvae were incubated in a petri dish with a small paper towel containing Suprane for 1–2 minutes in a fume hood. Anesthetized larvae were positioned ventral side up on a glass slide between spacers made by transparent tape, which prevented extreme compression of the larvae. Different size spacers were required for the various larval stages. Larvae were covered with a thin film of halocarbon oil and then with a cover glass. NMJ synapses on muscle 26 in hemi-segment 2 or 3 were imaged. After an imaging session, larvae were placed in numbered chambers with food in a 25°C incubator. The same data acquisition settings where used to visualize NMJs at different larval stages. Larvae were imaged with either 6, 24 and 36 hours intervals for one data set (Figure 8 A-C), or for 24 hours intervals for the remaining datasets. To keep the size consistent between different time periods, images of the corresponding NMJ area at younger stages were cut (dashed areas in figures) and placed onto a black background. This presentation generated a similar orientation of the different size NMJs for easier comparison for Figure 8, Figure 9 and Figure 8-supplemental figure 2.

### Statistical analysis

Statistical analysis was performed with GraphPad Prism using one-way ANOVA for comparison of samples within an experimental group, or Student’s t-test for comparing two groups. Asterisks denote p-values of: *, P≤0.05; **, P≤0.01; and ***, P≤0.001. All histograms and measurements are shown as mean ± SEM. Pearson coefficient of correlation was calculated in GraphPad Prism using the following parameters: - two-tailed P value and 95% confidence interval.

## Acknowledgements

This work was supported by NIH grant MH104536 to J.T.L. K.L.C. was supported in part by NIH pre-doctoral training grant T32GM007287. We thank the Bloomington *Drosophila* Stock Center (NIH P40OD018537), the Developmental Studies Hybridoma Bank, Richard Ordway (Penn State University) and Stephan Sigrist (Freie Univesitat Berlin) for providing *Drosophila* strains, Eliza Vasile (Koch Institute) for help with SIM data acquisition, and members of the Littleton lab for helpful discussions and comments on the manuscript.

## Supplemental Figure Legends

**Figure 4 - supplemental figure 1.**
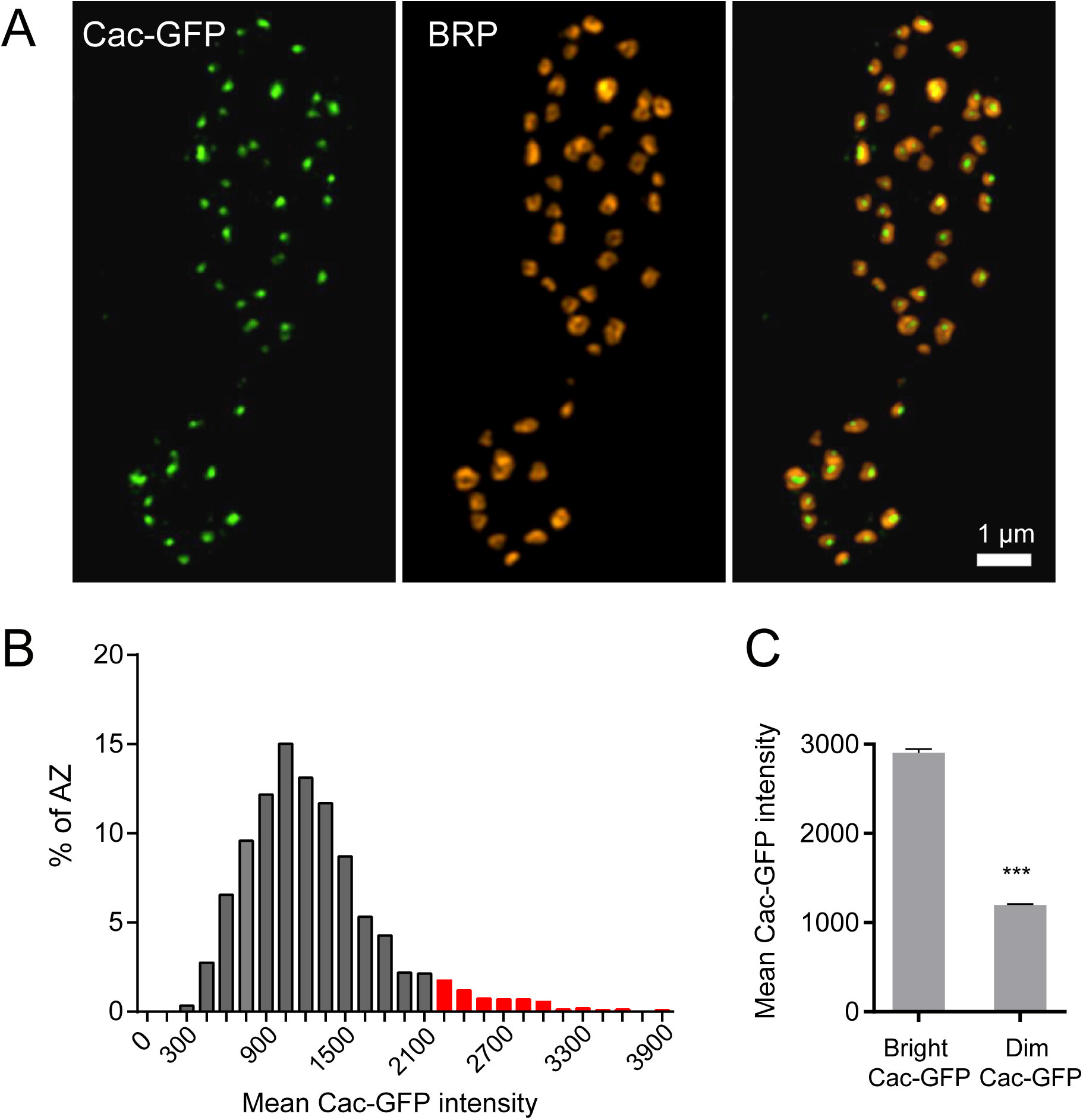
Cac-GFP distribution at AZs analyzed by SIM microscopy. (**A**) Representative Cac-GFP and BRP puncta at AZs for two synaptic boutons imaged using SIM microscopy. (**B**) Histogram of the distribution of mean Cac-GFP fluorescence intensity across the AZ population. Red corresponds to the Cac-GFP containing AZ population with fluorescence intensity 2 standard deviations above the mean. (**C**) Mean fluorescence intensity of Cac-GFP for bright (fluorescence greater than 2 standard deviations above the average) versus dim AZs. Student’s t-test was used for statistical analysis (*** = p≤0.001). Error bars represent SEM.

**Figure 7 - supplemental figure 1.**
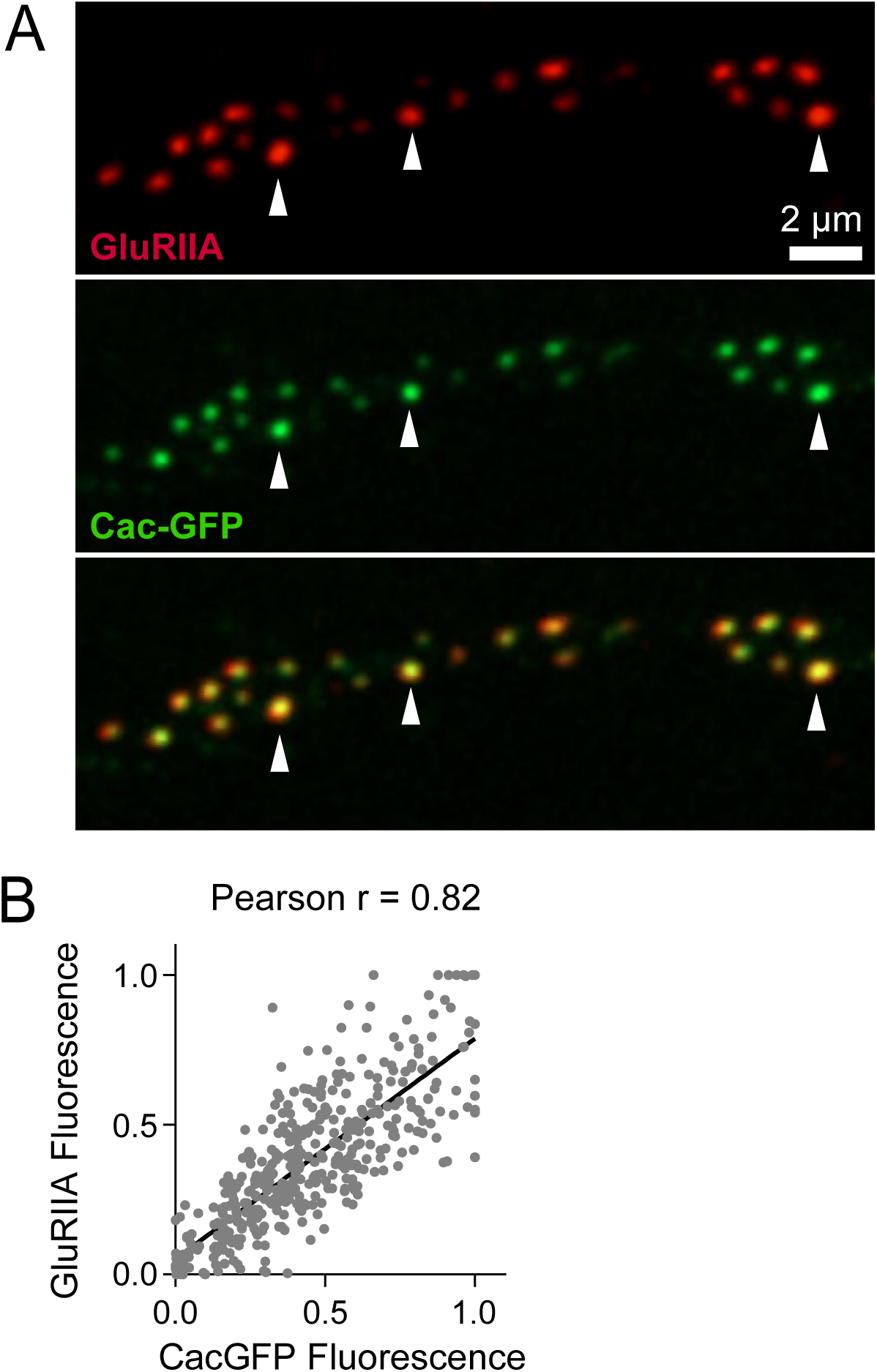
Correlation between GluRIIA and Cac fluorescence intensities during early larval development. (**A**) Representative Cac-GFP (green) and GluRIIA-RFP (red) synaptic puncta at a muscle 26 NMJ imaged through the cuticle of an anesthetized animal during early larval development. Arrows denote AZs with bright Cac-GFP opposed to PSDs with high levels of GluRIIA. (**B**) Correlation between GluRIIA-RFP and Cac-GFP at individual AZs.

**Figure 8 - supplemental figure 1.**
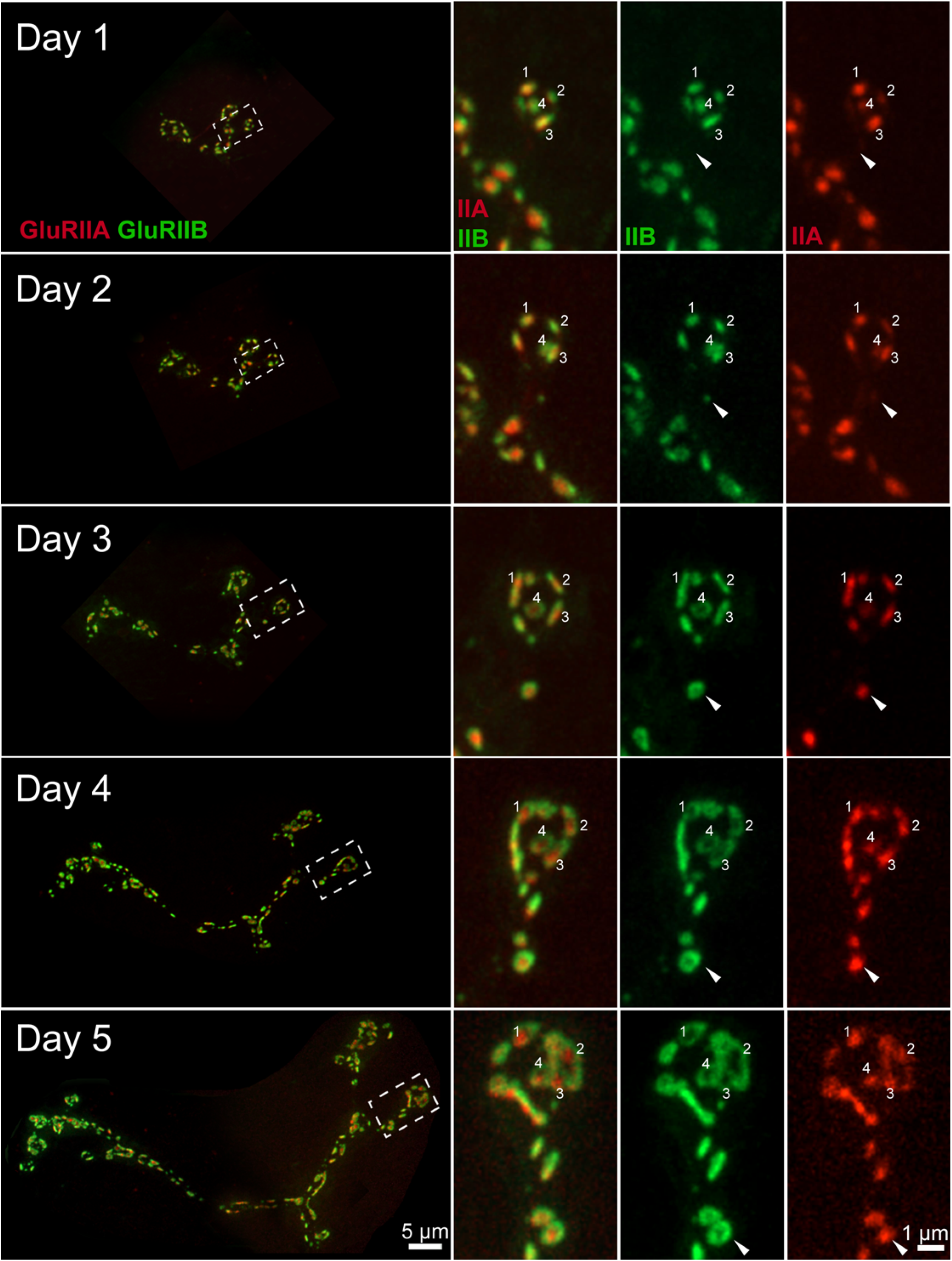
Consecutive imaging of NMJ growth at muscle 26 over a 5-day period imaged through the cuticle of an anesthetized larva during development. The entire NMJ is shown on the left panel. A smaller area (dashed box) is magnified and shown in the right panels. The merged image of GluRIIA and GluRIIB is shown in the middle, with individual GluRIIB and GluRIIA channels on the right. Several PSDs are numbered for tracking across imaging sessions. New PSDs appearing on day 2 (arrow) form the GluRIIB donut by day 4. On day 5, a larger number of PSDs display the characteristic GluRIIB peripheral segregation around a GluRIIA core, including those identified on the 1^st^ day of imaging (numbered).

**Figure 8 - supplemental figure 2.**
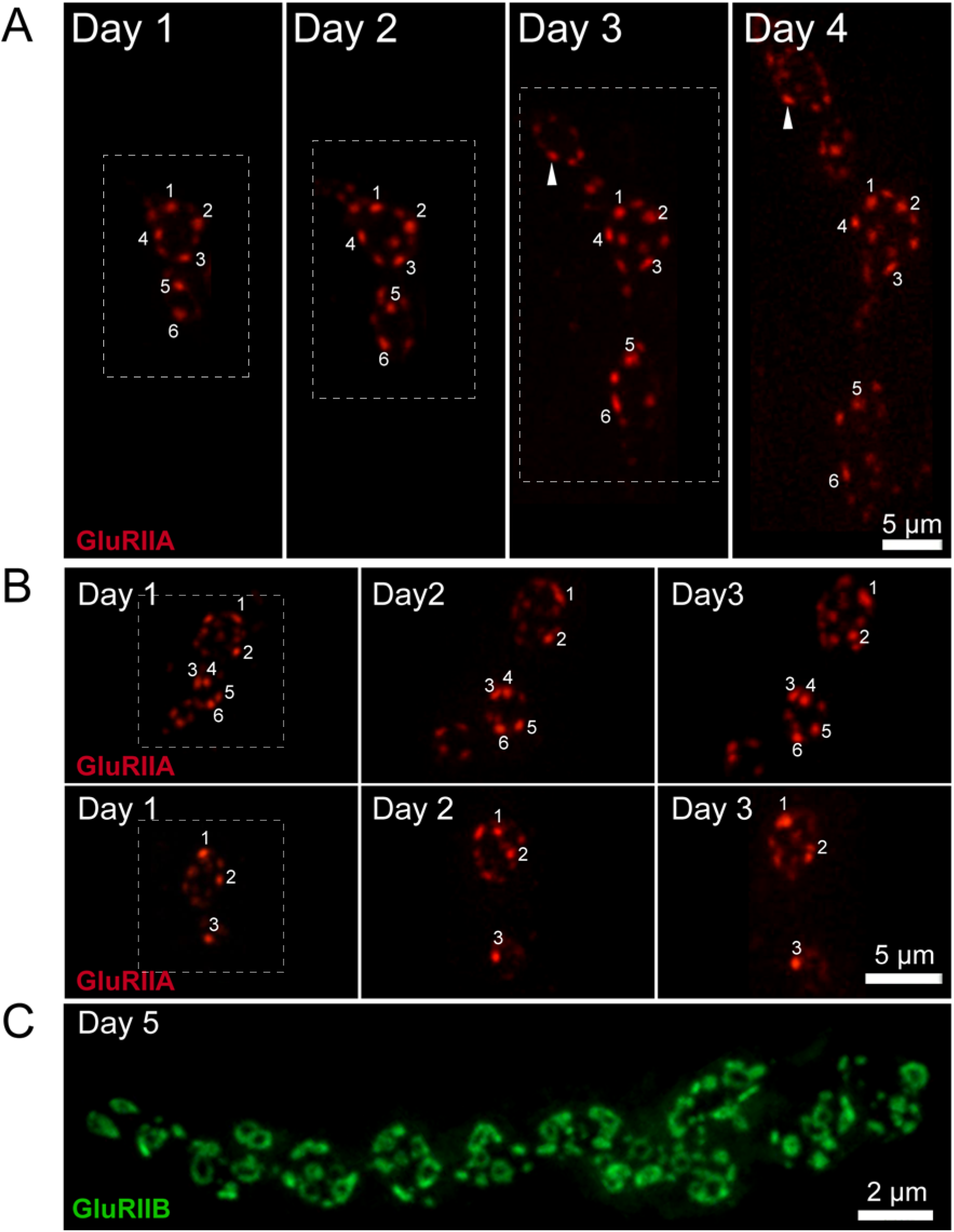
GluRIIA fluorescence intensity increases over time in a rate proportional to PSD birthdate. Representative serial images of muscle 26 NMJs visualized through the cuticle of two anesthetized larvae during development (**A**, **B**). The dashed box surrounds the actual imaged segment of the NMJ in each panel. The brightest GluRIIA puncta are numbered and followed through the imaging period. The brightest GluRIIA puncta observed on day 1 were among the brightest puncta on later days. Rarely, formation of new PSDs that showed a faster rate of GluRIIA accumulation were observed (arrow). (**C**) By day 5 of larval development, many PSDs show the donut-like GluRIIB distribution, though the number of smaller GluRIIB puncta also increase due to new AZ addition.

**Figure 8 - supplemental figure 3.**
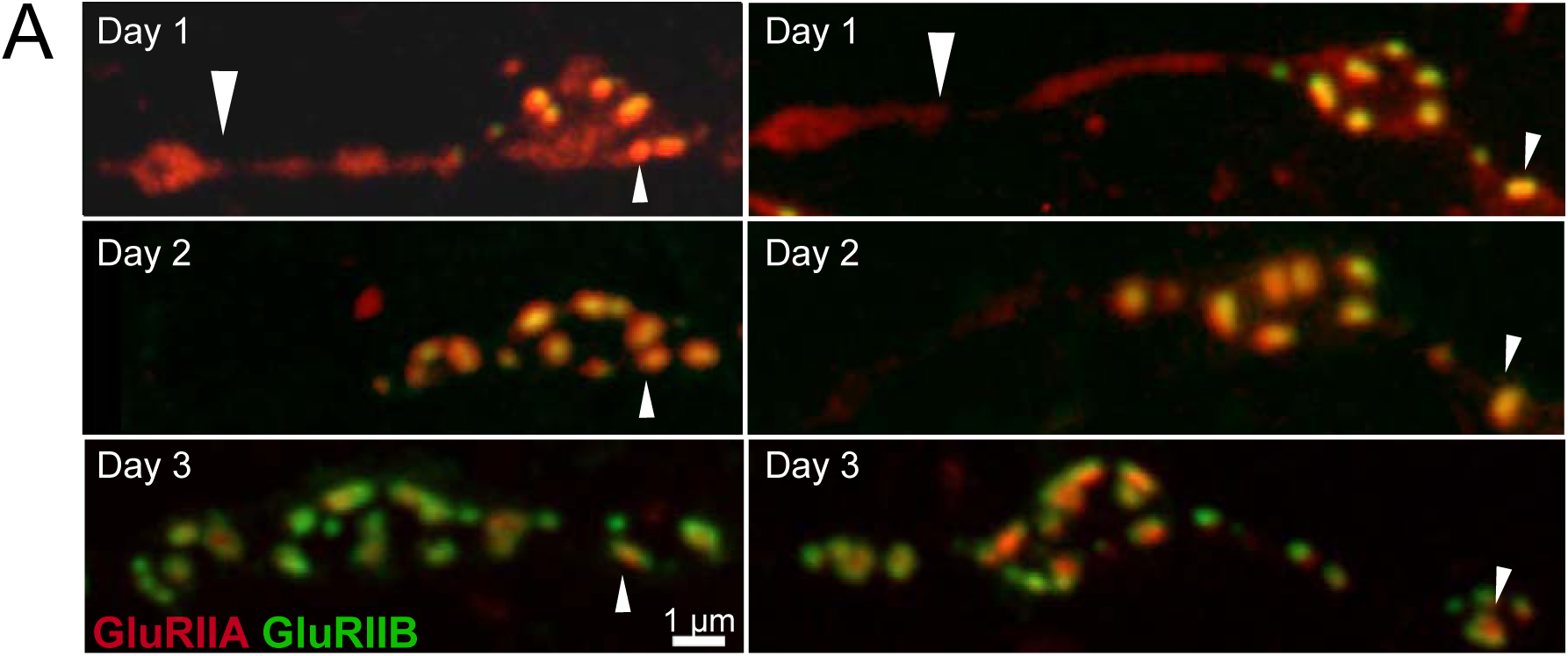
Synapse development along early GluRIIA positive NMJ extensions. (**A**) The two representative muscle 26 NMJs shown in Figure 8A were followed during development. The top panels display the NMJ structure visualized during live imaging in early 1^st^ instar larvae expressing GluRIIA-RFP and GluRIIB-GFP. The GluRIIA-RFP positive extensions from the main arbor that were devoid of detectable PSDs or GluRIIB at this stage of development (large arrow) later formed normal boutons with many PSDs by day 2 and 3. Smaller arrows denote the same PSD at each day for orientation.

**Movie 1**

Representative movie showing evoked and spontaneous GCaMP6s events (green) in larvae expressing GluRIIA-RFP (red) that were stimulated at 0.3 Hz.

**Movie 2**

Representative movie showing spontaneous GCaMP6s events in *syt1* mutants expressing GluRIIA-RFP (red), followed by GCaMP6s events observed during 5 Hz stimulation.

**Movie 3**

Representative movie showing evoked and spontaneous jRGECO events (red) in larvae expressing Cac-GFP (green).

